# Massively parallel characterization reveals context-dependent and non-additive regulatory effects of closely spaced variant pairs

**DOI:** 10.64898/2026.09.10.750584

**Authors:** Rita Kreevan, Vasili Pankratov, Danat Yermakovich, Navya Shukla, Mait Metspalu, Anders Eriksson, Michael Dannemann, Irene Gallego Romero, Tõnis Org

## Abstract

Precise control of gene expression relies in part on cis-regulatory elements (CREs), including enhancers. Genetic variation within enhancers can alter their regulatory activity and contribute to variation in gene expression, yet the effects of multiple nearby variants within the same enhancer remain poorly understood. To address this, we designed a massively parallel reporter assay (MPRA) to test the regulatory effect of 7,285 pairs of close-proximity single-nucleotide variants (SNVs) selected from blood-specific and broadly active enhancers. To enrich for functionality, we required at least one variant in each pair to be annotated as an eQTL in eQTLGen dataset. We measured the regulatory activity of all four haplotypes in K562 leukemia cells, where 57% of tested variant pairs had at least one derived haplotype that differed significantly in activity from the ancestral haplotype. The effects of individual variants frequently depended on the allelic background provided by the neighboring variant, with some variants showing opposite effects in different allelic backgrounds. Among a smaller, high-confidence subset selected for analysis of additivity, 59% (105/178) of variant pairs showed non-additive effects, and non-additive pairs were located closer together than additive pairs. Among pairs in which both single-derived haplotypes increased activity, the double-derived haplotype generally showed a smaller effect than expected under additivity. Together, our results demonstrate that nearby variants within the same enhancer can jointly shape regulatory activity and highlight the importance of considering local allelic context when interpreting the functional effects of regulatory variation.

## Introduction

Precise control of gene expression is essential for fundamental biological processes such as development, cellular differentiation, and the maintenance of cell identity. This control is mediated in part by cis-regulatory elements (CREs), including enhancers, which regulate the timing, location, and level of gene expression (Jindal & Farley, 2021; Long et al., 2016; Maston et al., 2006; Pennacchio et al., 2013; Sabarís et al., 2019). The regulatory activity of enhancers can be influenced by genetic variation within their sequences, thereby contributing to variation in gene expression levels (Chatterjee & Ahituv, 2017; Miguel-Escalada et al., 2015).

Genome-wide association studies (GWAS) have identified thousands of loci associated with complex traits and diseases. However, these associations generally do not identify the causal variants or reveal molecular mechanisms underlying their effects. Many trait-associated variants are located within CREs and are thought to exert their effects by altering gene regulation (Abdellaoui et al., 2023; Cano-Gamez & Trynka, 2020). Expression quantitative trait locus (eQTL) studies further support the link between genetic variation and gene expression by associating genetic variants with differences in gene expression levels (Brotman et al., 2025; Bryois et al., 2022; Nicolae et al., 2010; Võsa et al., 2021; Zivotic et al., 2025). At the molecular level, regulatory variants can alter CRE activity, for example by modifying transcription factor binding sites (TFBSs) and consequently transcription factor binding (Degtyareva et al., 2021; Deplancke et al., 2016; Dogan et al., 2015; Georgakopoulos-Soares et al., 2023; Grossman et al., 2017; Johnston et al., 2019; Rao et al., 2021). However, the mechanisms by which individual variants affect CRE activity and ultimately gene expression remain incompletely understood (Abdellaoui et al., 2022; Cano-Gamez & Trynka, 2020; Gallagher & Chen-Plotkin, 2018; Julienne et al., 2021).

Most studies of regulatory variation have focused on studying the effects of individual variants (Choi et al., 2020; Colli et al., 2021; Griesemer et al., 2021; Madan et al., 2019; Matoba et al., 2020; Tewhey et al., 2016; Ulirsch et al., 2016), yet multiple variants can occur in close proximity within the same enhancer and jointly influence its activity. Their combined effects may differ from those expected based on the effects of each variant individually. Rare variants are particularly relevant in this context, as variants with larger effects on gene expression tend to occur at lower frequencies, partly as a consequence of purifying selection (Li et al., 2017; Montgomery et al., 2011; Zhu et al., 2011). Together, these observations motivated us to investigate how nearby regulatory variants interact and how their effects relate to allele frequency. To address these questions, we used a haplotype-based massively parallel reporter assay (MPRA) to test all four allele combinations of closely spaced SNV pairs located within enhancers. This design allowed us to systematically characterize the individual and combined effects of these SNVs to determine how the regulatory effect of each variant is influenced by the neighbouring allelic background.

## Results

### Design of an MPRA library to assess the effects of closely spaced variant pairs

To construct the MPRA library, we selected naturally occurring bi-allelic SNV pairs from the Estonian Biobank (EstBB) whole-genome sequencing (WGS) data using several criteria (**Fig. 1a**). First, we required pairs to be separated by no more than 75 bp, so that both variants could be captured within a single MPRA oligonucleotide with some genomic context. Second, to enrich variants with regulatory effects, we prioritized pairs in which at least one variant was annotated as a cis-eQTLs in blood eQTLGen v1 (Võsa et al., 2021) (Methods). Third, because eQTLGen predominantly captures regulatory effects in blood, we focused on variant pairs located within blood-specific enhancers. We defined blood-specific enhancers as regions annotated as enhancers in at least 2 blood cell types, with at least one of them being a B-cell type, but not in non-hematopoietic cell types. For comparison, we also included variant pairs within broadly active enhancers, defined as regions annotated as enhancers in more than 85 tissues or cell types in the Roadmap Epigenomics (Roadmap Epigenomics Consortium et al., 2015) dataset (**Fig. 1b**). Finally, we designated one of the alleles as “ancestral” (Anc) and the other as “derived” (Der) based on the ancestral state reconstruction in human ancestral genome sequence from Ensemble (Dyer et al., 2025; Paten et al., 2008). These selection criteria resulted in 7,285 variant pairs of which 6,343 were located in blood specific enhancers and 942 in broadly active enhancers. For each variant pair, we designed four oligonucleotides, each representing one of the four possible haplotypes defined by the combinations of ancestral and derived alleles at the two variant positions (**Fig. 1c**). We refer to these four haplotypes collectively as a “haplotype set.” The final library comprised 29,140 haplotypes across 7,285 haplotype sets, and 860 control sequences, for a total of 30,000 sequences (Methods; Table S1). As our library was enriched for variants located within blood-specific enhancers, we performed seven replicate MPRA experiments in both the K562 (myelogenous leukemia) and GM12878 (lymphoblastoid) cell lines to assess their activity in two hematopoietic cellular contexts (**Fig. 1d**).

**Fig. 1.**
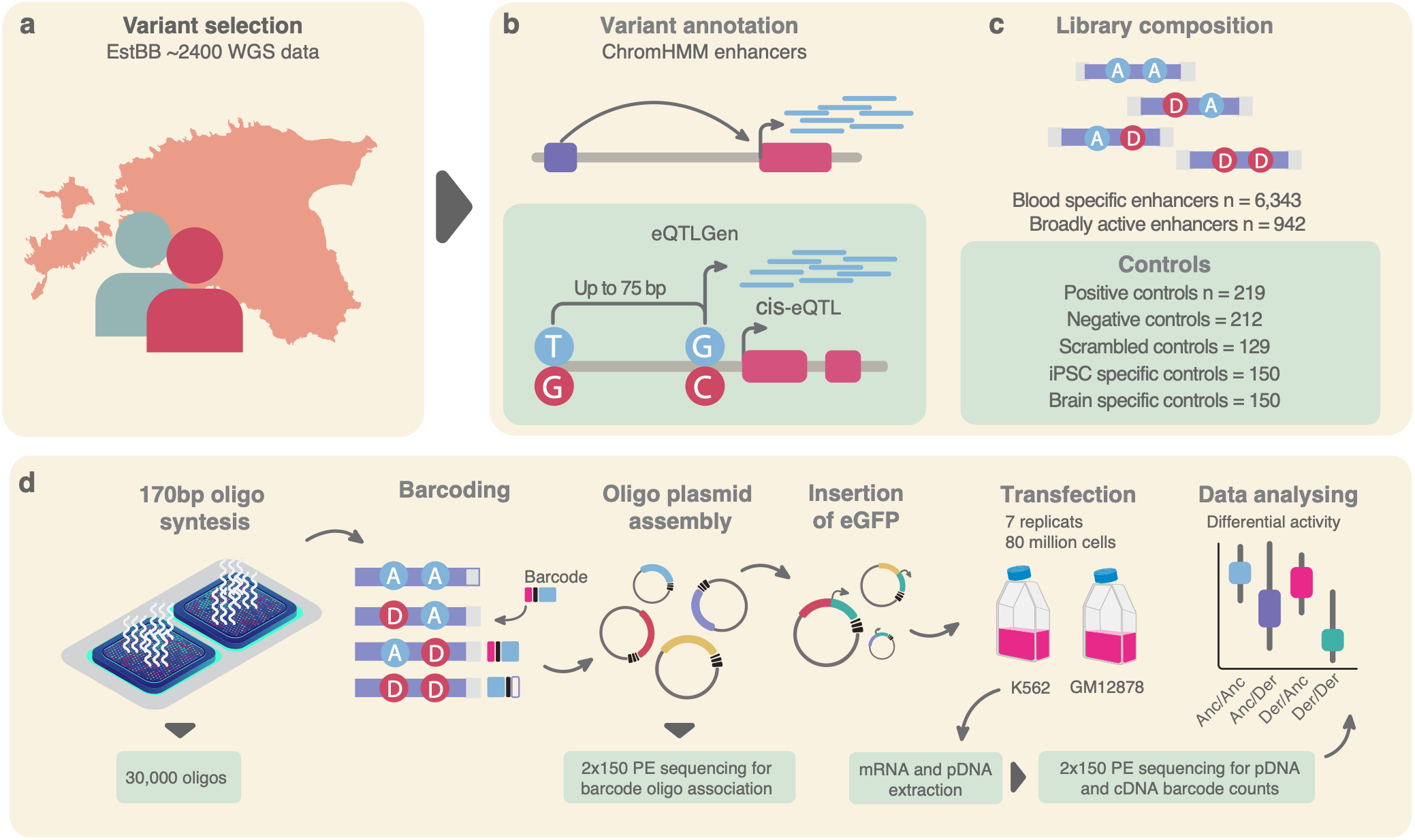
Overview of variant selection and MPRA workflow. **a** Variant pairs tested in the MPRA were selected from whole genome sequencing (WGS) data of ∼2,400 Estonian Biobank (EstBB) participants. **b** To prioritize variants with potential regulatory effects, variant pairs were annotated using Roadmap Epigenomics ChromHMM states. Blood-specific enhancers were defined based on enhancer annotations in B cells, while an additional set of broadly active enhancers was required to have enhancer annotations in >85 tissues and cell types. In addition, at least one variant in each pair was required to be a cis-eQTL in the eQTLGen dataset. **c** The final library contained 7,285 variant pairs, for which we designed all four possible haplotypes (allele combinations) together with 860 control sequences. **d** MPRA oligonucleotides (170 bp) were synthesized, PCR amplified, uniquely barcoded and cloning into plasmid backbone. The oligo-plasmid library was sequenced to determine variant-barcode associations. An eGFP open reading frame and minimal promoter were then inserted between the regulatory sequence and barcode by Gibson assembly to generate final MPRA library. The library was transfected into K562 and GM12878 cell lines. After 24h, RNA and DNA were extracted and sequenced, and barcode counts were used to quantify overall activity as the log_2_ RNA/DNA ratio. For each non-ancestral haplotype (Anc/Der, Der/Anc and Der/Der), differential activity (DA) was calculated as the difference in activity relative to the ancestral (Anc/Anc) haplotype.

Three GM12878 replicates (1, 5, and 7) were excluded from downstream analyses because cDNA abundance was highly correlated with input pDNA abundance (Pearson’s r = 0.83, 0.89, and 0.90, respectively; **Supplementary Fig. S2a, b; S3a, b**), indicating that transcript abundance largely reflected plasmid abundance rather than enhancer-driven transcription (Uebbing et al., 2021). Among the retained replicates, pDNA abundance was highly reproducible across replicates (Pearson’s r = 0.91-0.94), while cDNA abundance showed somewhat lower but still strong reproducibility (Pearson’s r = 0.79-0.86; Supplementary Figs. S2 and S3). Following data processing and quality control (Methods), we retained 5,988 haplotype sets in K562 and 6,106 in GM12878 cells for downstream analyses.

The overall activity of each tested haplotype was quantified as the log_2_ fold-change between RNA and DNA abundance across replicates (hereafter “activity”), using a negative binomial regression model implemented in DESeq2 (Love et al., 2014) (Methods). Within each haplotype set, we compared activity levels of the three haplotypes carrying one or more derived alleles against the ancestral/ancestral haplotype for evidence of differential activity (DA) between them. With positive and negative values indicating increased and decreased regulatory activity, respectively (Methods). Haplotypes with an adjusted *p*-value <0.01 after Benjamini–Hochberg correction were considered differentially active (Table S2; Table S3) (Methods).

To validate assay performance, we compared the activity of positive and negative control sequences with that reported for the same sequences by Tewhey et al. (Tewhey et al., 2016). Sequences that showed transcriptional activity in their assay were used as positive controls and sequences lacking detectable activity were used as negative controls. We then compared our activity measurements for these sequences to the results reported by Tewhey et al. As expected, positive controls showed higher median activity than negative controls in both tested cell lines (**Supplementary Fig. S4a, b, d, e, g**), consistent with the original study. Activity measurements for individual control sequences showed modest correlations between the two studies (Spearman’s *ρ* = 0.26, *p* = 1.35 x 10^-4^ for positive and *ρ* = 0.26, *p* = 4.05 x 10^-3^ for negative controls in K562; Spearman’s *ρ* = 0.32, *p* = 3.16 x 10^-6^ for positive and *ρ* = 0.32, *p* = 5.17 x 10^-4^ for negative controls in GM12878; **Supplementary Fig. S4a, 4d**). Similar differences between MPRA experiments have been reported previously (Jagoda et al., 2023) and may reflect differences in experimental design and data processing (Zhang et al., 2026). Overall, the clear separation between positive and negative controls supported the ability of our assay to distinguish sequences with different regulatory activities (**Supplementary Fig. S4c, f**).

We then asked whether the detected DA occurred in sequences with measurable transcriptional activity above background in both cell lines, using our set of (n=129) scrambled sequences as additional set of negative controls (Supplementary **Fig. S4b, c, e, f)**. As scrambled sequences should not drive transcription, their residual activity provides an estimate of background activity (Rosen et al., 2026). Scrambled controls had a median activity of 0.02, which we used as the background threshold (Supplementary **Fig. S4b, e**). Overall, 43% of the blood specific enhancers and 54% of broadly active enhancer DA haplotypes had activity above this threshold (Supplementary **Fig. S4c, f**), indicating that a substantial proportion of differential activity occurred in sequences with measurable transcriptional activity above background.

We next characterized DA in K562 and GM12878 cells. In K562, 57% (3,408/5,988) of haplotype sets contained at least one DA haplotype, compared with 14% (861/6,106) in GM12878. Despite the lower proportion of DA sets in GM12878, 77% (662/861) were also DA in K562 (**Fig. 2a; Table S3**). In total, 3,607 haplotype sets were DA in at least one cell line, including 2,746 unique to K562 and 199 unique to GM12878 (**Fig. 2a**). The number of shared DA sets was greater than expected by chance (Fisher’s exact test, OR = 2.93, 95% CI = 2.47-3.48, *p* = 2.02 x 10^-40^). The lower number of DA sets detected in GM12878 may partly reflect reduced statistical power following the exclusion of three of seven replicates during quality control, leaving the minimum number of replicates recommended for adequate power in MPRA experiments (Myint et al., 2019). We therefore performed the subsequent characterization of DA patterns and variant interactions in K562, for which all seven replicates were retained (**Table S4**).

**Fig. 2.**
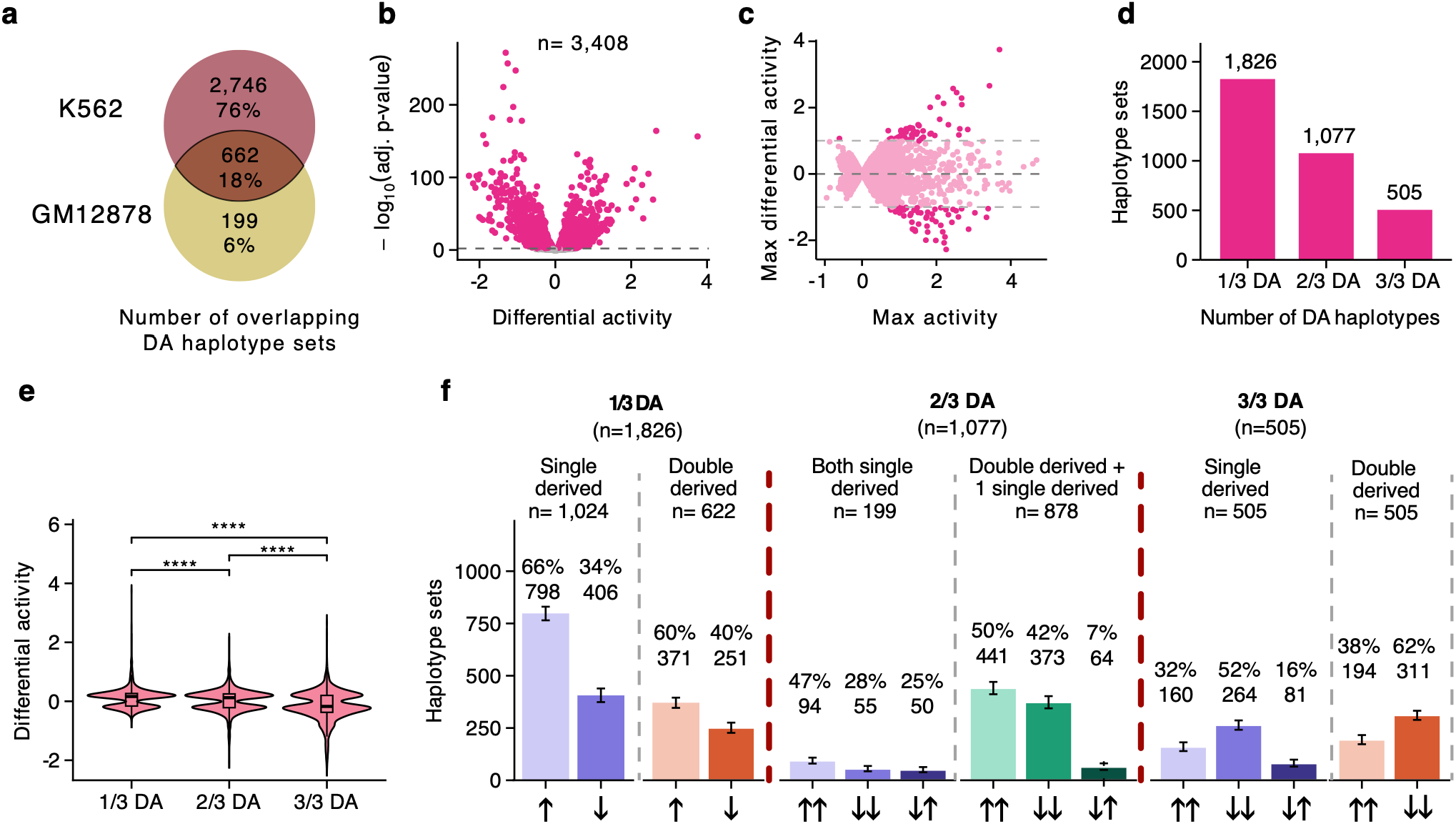
Identification of differentially active haplotypes. **a** Venn diagram showing an overlap of haplotype sets containing at least one differentially active (DA) haplotype between K562 and GM12878 cells. **b** Volcano plot showing differential activity of individual non-ancestral haplotypes in K562 cells. The dashed line indicates the significance threshold (Benjamini–Hochberg-adjusted *p* = 0.01). **c** Relationship between regulatory activity and DA across haplotype sets. The x-axis shows the maximum activity within each haplotype set and the y-axis the maximum DA relative to the ancestral haplotype (Anc/Anc). DA sets are shown in pink and non-DA sets in grey. Dashed lines indicate |DA| = 0.5. **d** Distribution of DA sets among the 1/3, 2/3, and 3/3 DA categories, defined by whether one, two, or all three non-ancestral haplotypes, respectively, were DA relative to Anc/Anc. **e** Violin plot showing the distribution of DA effect sizes across the three DA categories. **f** DA patterns within the 1/3, 2/3, and 3/3 DA categories. For the 1/3 and 2/3 DA categories, haplotype sets were classified according to which non-ancestral haplotypes were DA. In the 1/3 DA category, sets were classified as double-derived only, when Der/Der was the only DA haplotype, or single-derived only, when either Anc/Der or Der/Anc was the only DA haplotype. In the 2/3 DA category, sets were classified as double-derived + one single-derived, when Der/Der and one of the two single-derived haplotypes were DA, or both single-derived, when Anc/Der and Der/Anc were DA while Der/Der was not. In the 3/3 DA category, all three non-ancestral haplotypes were DA; therefore, DA direction is shown separately for the double-derived haplotype (Der/Der) and the single-derived haplotypes (Anc/Der and Der/Anc). Arrows indicate DA direction relative to the ancestral haplotype (Anc/Anc): ↑, increased activity; ↓, decreased activity. Paired arrows indicate the directions of two DA haplotypes, with ↑↑ and ↓↓ representing concordant effects and ↓↑ representing opposite effects. Numbers and percentages above the bars indicate the number and proportion of haplotype sets in each direction category.

### Regulatory differences between derived and ancestral sequences

Among the 3,408 haplotype sets containing at least one DA haplotype (hereafter referred to as DA sets) in K562 cells, we identified a total of 5,495 individual DA haplotypes (**Fig. 2b; Table S3**). Most DA haplotypes showed modest differences in activity relative to the ancestral haplotype, with only 162 showing more than two-fold difference (**Fig. 2c**). To further investigate this relationship, we classified DA sets according to the number of derived haplotypes (out of three) in the set that were DA relative to the ancestral haplotype, with 1/3 DA, 2/3 DA and 3/3 DA denoting one, two or all three derived haplotypes being DA, respectively (**Fig. 2d**). The magnitude and direction of DA differed across the three categories was assessed using pairwise Wilcoxon rank-sum tests with Benjamini–Hochberg correction (**Fig. 2e**). The 1/3 DA and 2/3 DA categories both showed modest positive effect directions (median DA 0.16 and 0.12, respectively), although their distributions differed significantly (adjusted *p* = 1.06 × 10⁻¹⁰). In contrast, the 3/3 DA category was shifted toward negative effects (median DA = −0.17) and differed significantly from both the 1/3 DA (adjusted *p* = 1.19 × 10⁻⁶¹) and 2/3 DA (adjusted *p* = 6.09 × 10⁻³⁰) categories. We next asked whether this shift was associated with the number of derived alleles carried by a haplotype. Although the number of derived alleles did not explain the differences between DA categories, within the 2/3 and 3/3 DA categories, DA associated with double-derived haplotypes (Der/Der) tended to be smaller than those observed for single-derived haplotypes (Anc/Der and Der/Anc) in the same category (Supplementary **Fig. S5b**). This trend was consistent across DA categories but did not reach statistical significance in 2/3 DA and 3/3 DA haplotype sets (Wilcoxon rank sum test, 1/3 DA adj.p = 0.023; 2/3 DA adj.p = 0.334; 3/3 DA adj.p = 0.334; Supplementary Fig. S5b).

We then examined which haplotypes were DA within each haplotype set category, asking whether the regulatory effect associated with a focal derived allele was maintained when the allelic state of the other variant in the set changed. Because the non-ancestral haplotypes carry either one derived allele (Anc/Der or Der/Anc) or two derived alleles (Der/Der), their DA patterns allowed us to distinguish different allele combination effects (**Fig. 2f**; Supplementary **Fig. S5a**). In the 1/3 DA category, all three haplotypes appear to be equally likely to be DA relative to the Anc/Anc state: in 65.9% (1,204/1,826) the DA haplotype has one derived allele (Der/Anc or Anc/Der) and two derived alleles (Der/Der) in 34.1% (622/1,826) (**Fig. 2f**; Supplementary **Fig. S5a**). Overall, activity-increasing effects were more common than activity-decreasing effects among 1/3 DA sets (64%; 1,169/1,826 vs. 36%; 657/1,826; binomial test, *p* < 2.2 × 10⁻¹⁶; **Fig. 2f**). These patterns indicate that the regulatory effect associated with a derived allele can depend on the allelic state of the neighboring variant.

In contrast, in the 2/3 DA category having one common derived allele across the two DA haplotypes was more common than expected by chance: this occurred in 81.5% (878/1,077) of sets compared to the expected 66.6%. In most of these sets (92.7%; 814/878), the two DA haplotypes showed effects in the same direction, whereas in 7.3% (64/878) they had opposite effects (**Fig. 2f**; Supplementary **Fig. S5a**). In the remaining 18.5% of 2/3 DA sets (199/1,077), where there was no common derived allele across the two DA haplotypes, the DA haplotypes showed effects in the same direction in 74.9% (149/199) of these sets and in opposite directions in 25% (50/199) (**Fig. 2f**; Supplementary **Fig. S5a**). Thus, although most 2/3 DA sets showed a consistent effect associated with one derived allele across two haplotypes, a subset again showed DA patterns that were dependent on the allelic state of the neighboring variant.

The 3/3 DA category represents a more complex pattern in which both single-derived haplotypes (Anc/Der and Der/Anc) and the double-derived haplotype (Der/Der) were DA relative to the ancestral haplotype (**Fig. 2f**). Among the 505 3/3 DA sets, the two haplotypes containing a single derived and ancestral alleles showed effects in the same direction in 424 sets (84%), with both increasing activity in 160 sets and, both decreasing activity in 264 sets. The DA haplotypes with two derived alleles showed a bias toward decreased activity (311 vs. 194 sets). Furthermore, all three non-ancestral haplotypes decreased activity in 52% of all 3/3 DA sets (263/505). When looking at only those haplotype sets where all three haplotypes had the same DA direction, they were significantly more likely to decrease activity rather than increase (263 vs. 158 sets; binomial test, *p* = 3.5 x 10^-7^; **Fig. 2f**).

### Variant effects are often dependent on the neighboring variant

Among the 5,988 haplotype sets tested in K562 cells, 1,426 variants were included in two different haplotype sets, each paired with a different neighboring variant (**Fig. 3a**). This allowed us to assess whether the regulatory effect of a genetic variant was maintained across different neighboring genomic backgrounds. Of these variants, 37.6% were DA in one background but not the other (**Fig. 3b, c, g; Supplementary Fig. S6c-d**), and their effects showed no strong correlation between backgrounds (Pearson’s r = 0.071, *p* = 0.1). In contrast, 11% (157/1,426) were DA in both backgrounds, and their effect sizes were strongly correlated (Pearson’s r = 0.61, *p* = 2.95 × 10⁻¹⁷; **Fig. 3b-f; Supplementary Fig. S6a-b**). Among these 157 variants, the direction of DA was maintained across backgrounds for 77.7% (122/157), whereas 22.3% (35/157) showed opposite directions (**Fig. 3e-f**). Variants maintaining the same direction between backgrounds did not show a significantly larger DA magnitude than those with opposite DA between backgrounds (Wilcoxon rank sum test, *p* = 0.24 and *p* = 0.23 for the two backgrounds; Supplementary **Fig. S6e**). Modest sample size for the opposite DA sets (n=35) limits the power to detect subtle differences. Moreover, the DA category (1/3, 2/3, or 3/3 DA) changed between backgrounds for 61% (96/157) of these variants (**Fig. 3d**). The remaining 51.4% (733/1,426) showed no DA in either background (**Fig. 3b-c**). Together, these results again show that the regulatory effects of nearby variants frequently depend on the variant background in which they are tested.

**Fig. 3.**
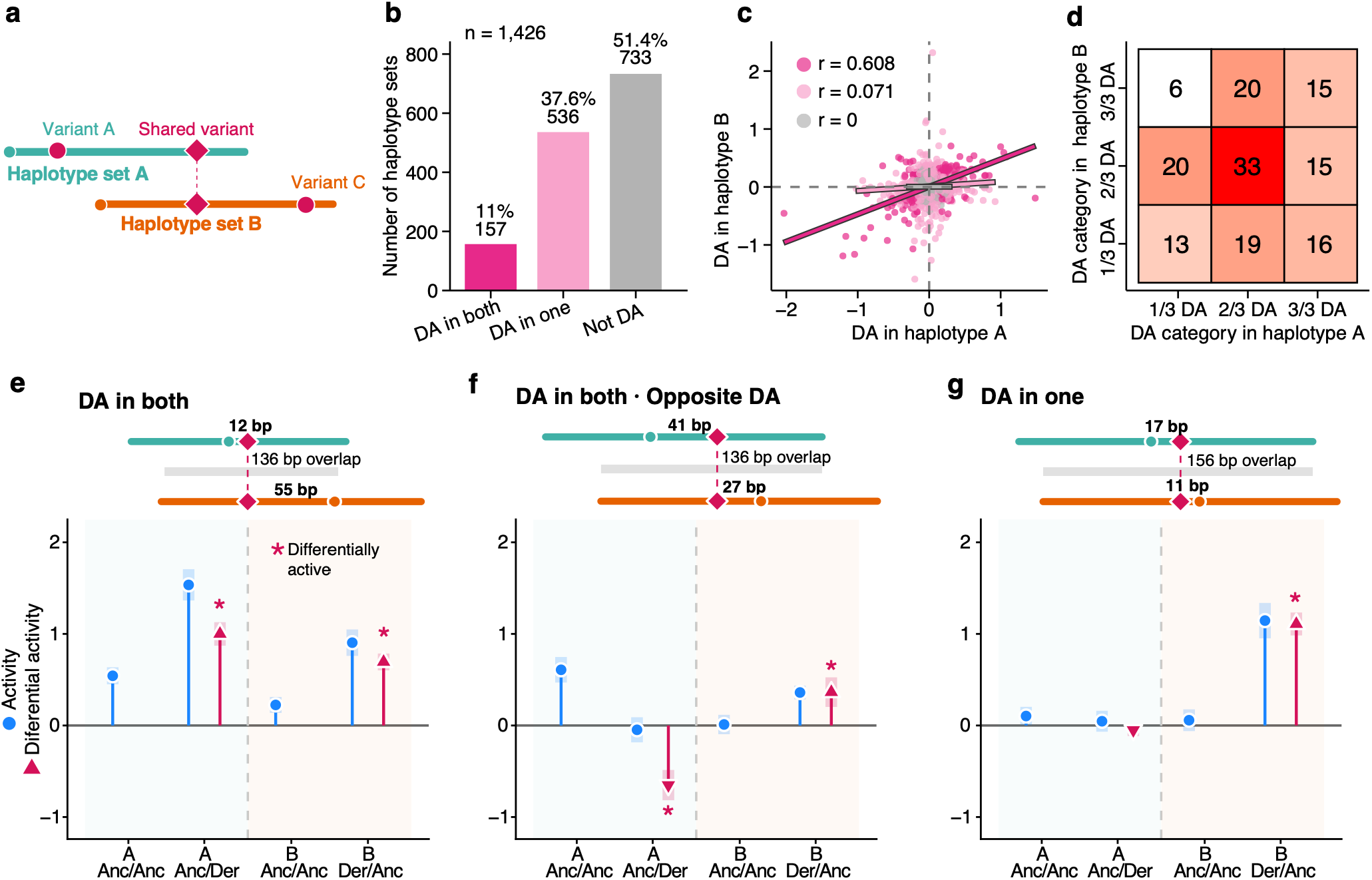
Regulatory effects of variants depend on the neighboring variant background. **a** Schematic illustrating the same variant tested in two different haplotype sets, each paired with a different neighboring variant. **b** Classification of variants tested in two haplotype sets according to their DA status across the two neighboring variant backgrounds: DA in both, DA in one, or not DA in either set. **c** Comparison of DA effect sizes for the same variant across the two haplotype sets. Variants are grouped according to whether they were DA in both sets (pink), DA in only one set (light pink), or not DA in either set (grey). **d** Comparison of DA categories across the two haplotype sets, showing whether the DA category was maintained or changed between neighboring variant backgrounds. **e** Example of a variant that was DA in both haplotype sets and showed the same direction of DA in both backgrounds. **f** Example of a variant that was DA in both haplotype sets but showed opposite directions of DA between the two backgrounds. **g** Example of a variant that was DA in only one of the two haplotype sets. In **e–g**, **overall** activity is shown in blue and DA relative to the ancestral haplotype in red; asterisks indicate haplotypes used to assess the effect of the shared variant.

### Genomic features associated with differential activity

We next examined whether DA was associated with genomic features of the tested variant pairs. We first assessed whether the occurrence or magnitude of DA varied with distance to the nearest transcription start site (TSS). The proportion of DA haplotype sets was similar across TSS distance bins (55–61%; Supplementary **Fig. S7a**), and the magnitude of DA did not differ significantly with TSS distance (ANOVA, *p* = 0.16; **Fig. 4a**). We then asked whether the distance between the two variants influenced DA. Similarly, the proportion of DA haplotype sets was similar across variant-distance bins (Supplementary **Fig. S7b**), and DA magnitude showed no significant association with variant distance (ANOVA, *p* = 0.16; **Fig. 4b**). Thus, within the genomic distances represented in our library, neither proximity to a TSS nor distance between paired variants was strongly associated with DA.

**Fig. 4.**
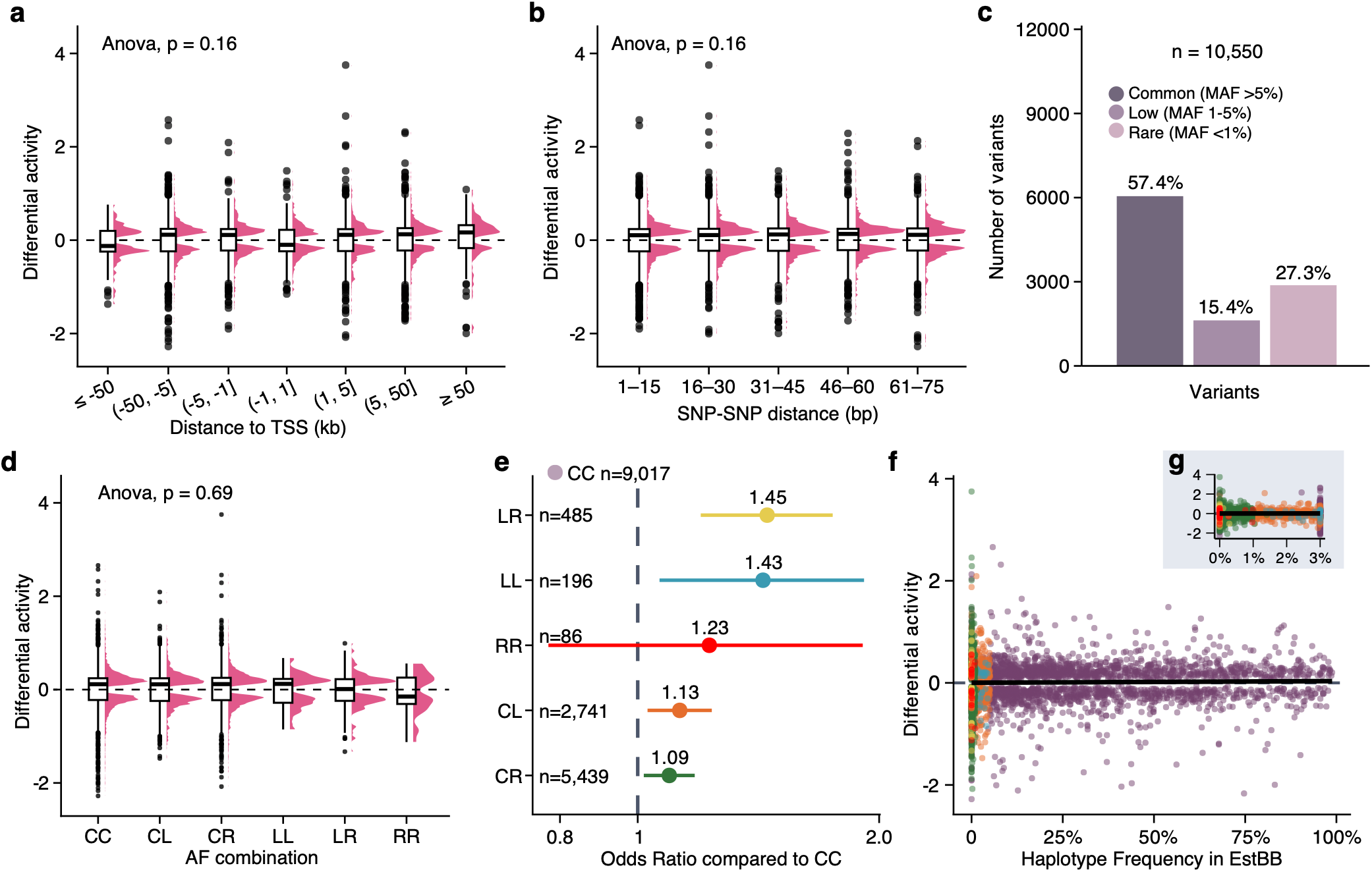
Genomic and population features associated with differential activity. **a** Distribution of DA effect sizes across bins defined by distance to the nearest transcription start site (TSS). **b** Distribution of DA effect sizes across bins defined by the distance between the two variants in each pair. **c** Minor allele frequency (MAF) distribution of variants in the EstBB, classified as common (MAF >5%), low-frequency (MAF 1–5%), or rare (MAF <1%). **d** Distribution of DA effect sizes across allele-frequency (AF) combinations. CC, common/common; CL, common/low-frequency; CR, common/rare; LL, low-frequency/low-frequency; LR, low-frequency/rare; RR, rare/rare. **e** Odds ratios for haplotypes being DA across AF combinations, relative to the common/common group. Numbers indicate the total number of haplotypes in each group. **f** Relationship between DA effect size and haplotype frequency in EstBB, with haplotypes colored according to their AF combination. **g** Enlarged view of **f** showing haplotypes with frequencies up to 3%.

### Allele frequency predicts differential activity but not its magnitude

We next examined whether allele frequency (AF), estimated from the Estonian BioBank cohort (n > 210,000 participants; 2,420 WGS), was associated with the occurrence or magnitude of DA. Approximately one quarter of the tested variants were rare (MAF <1%), 15% were low frequency (MAF 1–5%), and the remainder 57% were common (MAF >5%; **Fig. 4c**; Supplementary **Fig. S7c-e**). We classified variant pairs according to the AF categories of the alleles present in each haplotype and compared DA across these combinations. The magnitude of DA did not differ significantly among AF combinations (ANOVA, *p* = 0.69; **Fig. 4d**; Supplementary **Fig. S7c**), indicating that haplotypes containing rare alleles did not generally show stronger regulatory effects. In contrast, the odds of being DA slightly increased with lower AF (logistic regression, OR = 1.07 per frequency category; 95% CI 1.04-1.10, *p* < 1.51 × 10⁻^5^), with low-frequency/rare combinations showing up to ∼1.45-fold higher odds of DA than common/common combinations (**Fig. 4e**). Thus, rare alleles were slightly more likely to be DA, but their effects were not larger when they were classified as DA.

We next examined whether the population frequency of the haplotypes in EstBB was associated with DA. Of the 23,952 tested haplotypes, 9,722 had a frequency below 1% in the 2,420 EstBB WGS samples (Supplementary **Fig. S7f)**. These rare haplotypes were enriched among DA haplotypes (χ² test, *p* = 0.002) though with a modest effect size (OR ≈ 1.11). However, among haplotypes classified as DA, again, neither the magnitude nor direction of DA showed a clear association with haplotype frequency (**Fig. 4f**, **g**). Thus, rare haplotypes were more likely to be DA, whereas among DA haplotypes, neither effect magnitude nor direction was associated with haplotype frequency.

### Regulatory variants within a haplotype frequently interact non-additively, with effects shaped by proximity

The presence of two variants within each haplotype set allowed us to search for non-additive interactions between nearby regulatory variants. To reliably assess such interactions, we focused on a high-confidence (HC) subset of haplotype sets with substantial regulatory activity and DA effects. We required at least one of the four haplotypes in a set to have an absolute RNA/DNA activity log2FC >1 and at least one non-ancestral haplotype (Anc/Der, Der/Anc, or Der/Der) to have an absolute DA > 0.5 relative to the Anc/Anc haplotype. Applying these criteria retained 178 HC haplotype sets, comprising 712 individual haplotypes (**Fig. 5a**). The HC set contained a higher proportion of haplotype sets in which multiple haplotypes were DA (82%) and fewer haplotype sets with single DA haplotype (18%). Because HC sets were selected based on the presence of at least one non-ancestral haplotype with an absolute DA >0.5, the DA distribution within individual DA category differed from those of the full dataset (**Fig. 5b**). Compared with the full dataset, the HC subset showed a shift toward stronger positive effects in the 1/3 DA category and broader distributions of DA effect sizes in the 2/3 and 3/3 DA categories (**Fig. 5b**). To assess whether the strongest MPRA signals were also supported by potential regulatory activity in their endogenous genomic context, we examined their overlap with open chromatin regions identified by ATAC-seq and DNase hypersensitivity data in K562 cells. We ranked DA haplotypes and found that nearly all of the highest-ranking haplotypes overlapped open chromatin regions, with the proportion decreasing as progressively lower-ranking haplotypes were included (**Fig. 5c**). This enrichment provides independent support for the regulatory activity of regions showing the strongest effects in our MPRA.

**Fig. 5.**
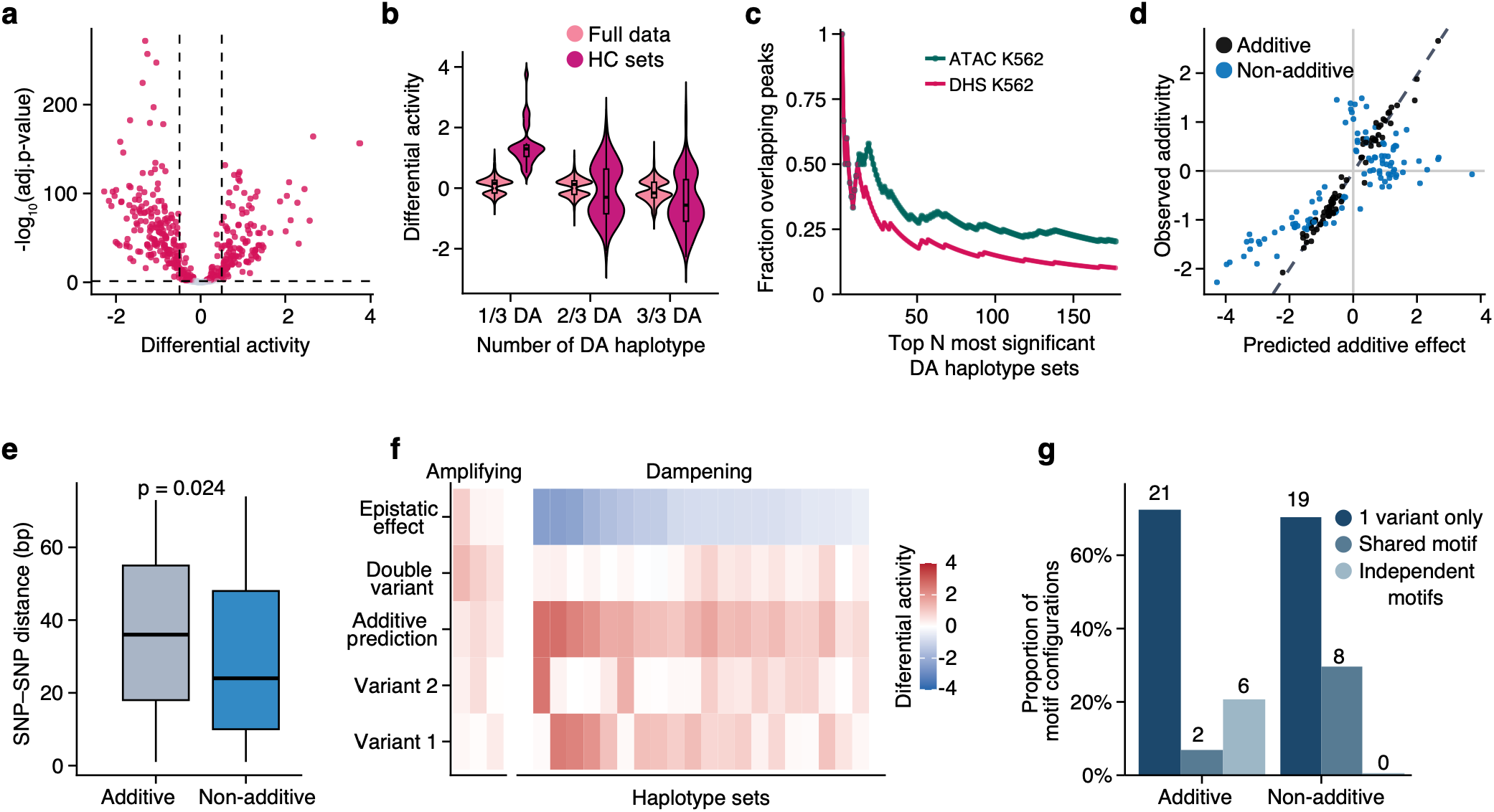
Non-additive regulatory interactions among high-confidence differentially active haplotype sets. **a** Volcano plot showing DA effect sizes and adjusted *p*-values for haplotypes within the high-confidence (HC) sets. Dashed lines indicate |DA| = 0.5. **b** Distribution of DA effect sizes among the 1/3, 2/3, and 3/3 DA categories within the HC sets. **c** Fraction of DA haplotypes overlapping open chromatin regions in K562 cells. Haplotypes were ranked by adjusted *P*-value for DA from most to least significant. **d** Classification of HC haplotype sets as additive or non-additive based on the relationship between the observed DA of the double-derived haplotype and the expected additive effect of the two single-derived haplotypes. **e** Genomic distance between the two variants in additive and non-additive haplotype sets. **f** Comparison of observed and expected DA in non-additive sets in which both single-derived haplotypes (Anc/Der and Der/Anc) showed activity-increasing effects. **g** TF motif-overlap configurations in additive and non-additive haplotype sets. 1 variant only indicates sets in which only one variant overlaps a predicted TF motif; Shared motif indicates sets in which both variants overlap the same predicted motif; Independent motifs indicates sets in which the two variants overlap separate predicted TF motifs.

We next tested whether the combined effects of the two variants deviated from additivity. For each haplotype set, we defined the expected additive effect as the sum of the DA of the two single-derived haplotypes (Anc/Der + Der/Anc). A haplotype set was classified as non-additive when the 95% CI of the observed DA of the double-derived haplotype (Der/Der) did not overlap with the 95% CI of this additive expectation, thereby ensuring the deviation exceeded the combined uncertainty of both estimates. Using this criterion, 59% (105/178) of HC haplotype sets showed non-additive effects (**Fig. 5d; Table S5**), indicating that the combined effects of nearby regulatory variants frequently deviated from additivity.

We then asked how the effects of the two individual variants combined in non-additive sets. Non-additivity was observed both when the two single-derived haplotypes had effects in opposite directions and when their effect directions were concordant. Focusing on HC sets in which both single-derived haplotypes independently increased activity, we found that the combined effect was generally smaller than expected from the sum of their individual effects (**Fig. 5f**). This predominantly sub-additive pattern is consistent with previous observations of regulatory epistasis (Siraj et al., 2026). Because close-proximity variants may more likely affect the same or interacting regulatory elements, we next asked whether non-additivity was associated with the distance between the two variants. Within the HC set, non-additive variant pairs were located closer together than variants in additive pairs (median 24 vs. 36 bp; Wilcoxon test, *p* = 0.024; **Fig. 5e**) consistent with previous findings (Siraj et al., 2026). This association supports a role for local sequence organization in shaping interactions between nearby regulatory variants. Together, these results show that nearby regulatory variants frequently interact non-additively, with their combined effects often being smaller than predicted from their individual effects.

To determine whether haplotypes showing non-additive regulatory effects are observed in the Estonian population, we examined the frequencies of the four possible haplotypes in each HC non-additive set (105 haplotype sets; 420 individual haplotypes) using EstBB WGS data. For most non-additive haplotype sets (96/105), three of the four possible haplotypes were observed in EstBB, whereas all four were observed for only three sets and only two for six sets (**Supplementary Fig. S8a**); 108 individual haplotypes were not observed in EsBB data at all. Notably, 3 of these missing haplotypes were ancestral, which we removed from the downstream comparison. Overall, 76.2% (80/105) of the haplotypes that were not observed were DA. Double-derived haplotypes were the most frequently absent haplotypes (67/105) and accounted for 53 of these 80 DA haplotypes. However, among the absent haplotypes, double-derived haplotypes were not significantly more likely to be DA than other haplotypes, not in the full dataset or within non-additive sets (OR = 1.54, CI 95% 0.55-4.22, *p* = 0.35; Supplementary **Fig. S8b-d)**. Thus, while haplotypes not observed in EstBB were likely to be DA and carry both derived alleles, this association was not significant. This likely reflects limited power given the small number of absent haplotypes. This pattern is broadly consistent with the possibility that regulatory effects deviating form an optimal level are kept at low frequencies in the population. Although the absence of these haplotypes from EstBB WGS data could also reflect their recent origin of limited power to detect very rare haplotypes.

### Additive and non-additive variants show similar motif density but distinct motif architecture

Transcription factor (TF) binding is a key mechanism through which regulatory variants can alter enhancer activity (Lambert et al., 2018; Spitz & Furlong, 2012). We therefore examined whether TF motif patterns differed between additive and non-additive HC sets. Because multiple TF motifs can occur within the 170-bp sequences tested in our MPRA, we first asked whether additive and non-additive sets differed in overall motif density. Of the 178 HC haplotype sets, 109 contained at least one known TF motif, on average two and up to 19 motifs per set (**Supplementary Fig. S8e**). The proportion of sets containing at least one motif did not differ between additive and non-additive sets (Fisher’s exact test, *p* = 0.35), nor did the total number of unique motifs per set (Wilcoxon test, *p* = 0.22; **Supplementary Fig. S8e**). Thus, overall motif content was similar between additive and non-additive sets.

We next asked whether the assayed variants themselves overlapped TF motifs, providing a more direct indication that sequence variation could alter a potential TF binding site. At least one variant overlapped a TF motif in 37% of additive sets (27/73) and 25.7% of non-additive sets (27/105), although this difference was not statistically significant (Fisher’s exact test, *p* = 0.074; **Supplementary Fig. S8f**). However, the number of unique motifs overlapping the variants was higher in additive than non-additive sets (Wilcoxon test, *p* = 0.043; **Supplementary Fig. S8g**). Thus, although the likelihood of having at least one variant motif overlap was similar between the two groups, additive sets tended to have more motifs overlapping the tested variants.

Given this difference, we asked whether additive and non-additive sets also differed in the arrangement of motif overlaps between the two variants. We classified these overlapping according to whether a motif overlapped only one variant, was shared by both variants, or whether the two variants overlapped distinct motifs. The distribution of these patterns differed significantly between additive and non-additive sets (Fisher’s exact test, *p* = 0.023; **Fig. 5g**). Additive sets showed a broader range of motif architectures, including 21 instances with a single-variant overlap, two of a shared motif, and six in which both variants overlapped distinct motifs, either alone or together with a shared motif. In contrast, non-additive sets included 19 single-variant overlaps and eight shared-motif cases, but no cases in which the two variants overlapped distinct motifs (**Fig. 5g**). These results suggest that non-additive interactions are preferentially associated with variants affecting a single or shared regulatory motif rather than two distinct motifs. Together with the shorter distances observed between non-additive variant pairs, this pattern suggests that local regulatory architecture may contribute to non-additive interactions between nearby variants.

### Motif contexts at trait-associated loci illustrate non-additive effects

To examine whether the non-additive effects identified in our MPRA occur at loci relevant to human traits, we assessed known GWAS (Cerezo et al., 2025; Sollis et al., 2023) associations among the variants within the 105 HC non-additive sets. Thirteen sets contained at least one genome-wide significant GWAS-associated variant (*P* < 5 x 10^-8^), with associations reported for traits including blood protein and circulating biomarker levels, hematological traits, immune-mediated diseases, cardiovascular disease, and psychiatric traits (Supplementary **Fig. S9**; **Table S6**). One example involved two adjacent variants within a blood-specific enhancer on chromosome 17: rs35689157 (chr17:37962845, G>A; MAF = 0.064 in EstBB) and rs77232433 (chr17:37962846, C>T; MAF = 0.064 in EstBB). In our MPRA, the ancestral haplotype showed the highest regulatory activity, whereas introduction of either derived allele reduced activity, with little additional effect when both derived alleles were present (**Fig. 6a**). The variants were in complete LD in EstBB (R² = 1, D′ = 1), such that only the ancestral haplotype (frequency in EstBB ∼94%) and double-derived haplotype (frequency in EstBB ∼6%) were observed (**Fig. 6b**). Both variants overlap the same predicted ZNF184 motif (**Fig. 6c**), suggesting that disruption of this shared motif by either substitution may be sufficient to reduce enhancer activity and thereby explain the non-additive effect. In eQTLGen, the derived alleles were associated with increased expression of several genes (*GSDMA, RP11-94L15.2*), including the strongest linked gene, *IKZF3*, but with decreased expression of *GSDMB* and *PGAP3*. Given the role of *IKZF3* in B-cell differentiation and the association of this locus with eosinophil count, this variant pair provides a trait-relevant example of non-additive regulatory effects occurring within a shared motif context.

**Fig. 6.**
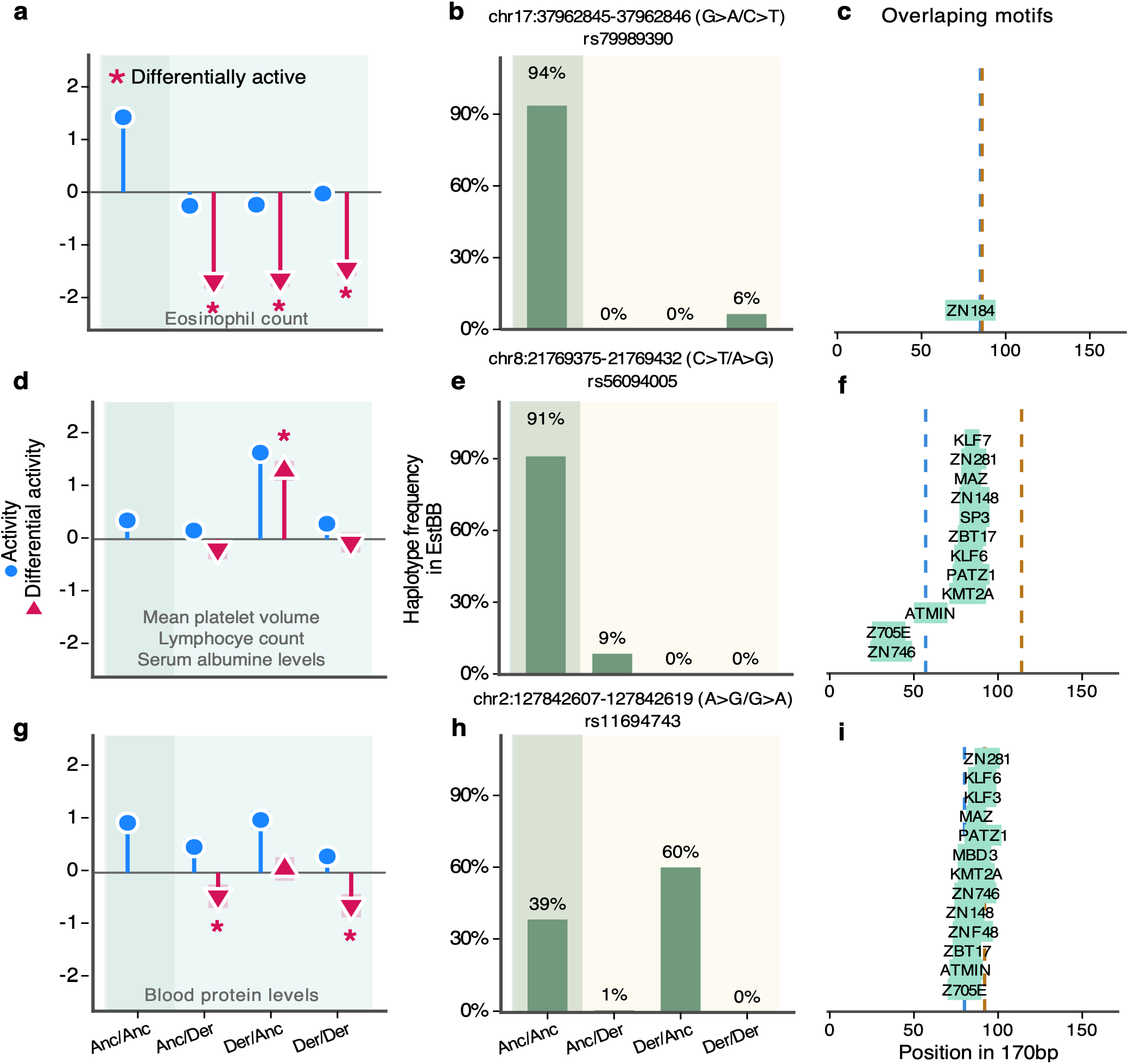
Examples of non-additive regulatory effects at loci with known GWAS associations. **a–c**, variant pair rs35689157 (chr17:37962845, G>A)–rs77232433 (chr17:37962846, C>T). **d–f**, variant pair rs373143737 (chr8:21769375, C>T)– rs56094005 (chr8:21769432, A>G). **g–i**, variant pair rs11694743 (chr2:127842607, A>G)–rs549250007 (chr2:127842619, G>A). For each variant pair, panels show MPRA regulatory activity and DA (**a, d, g**), haplotype frequencies in EstBB (**b, e, h**), and predicted TF motif overlaps within the 170-bp assayed sequence (**c, f, i**). Dashed lines in the motif panels indicate the positions of the two variants.

The remaining two examples illustrate non-additive effects in motif-rich sequence contexts, but with distinct motif architectures and population haplotype structures. The first variant pair, rs373143737 ( chr8:21769375, C>T) and rs56094005 (chr8:21769432, A>G) are located on chromosome 8 and are separated by 57 bp (**Fig. 6d**). The common variant, rs56094005 (A>G, MAF = 0.087 in EstBB; **Fig. 6e**) is an eQTL linked to *GFRA2*, *DOK2* and *NUDT18*. This variant is also associated with multiple hematological traits like platelet distribution width, mean platelet volume and lymphocyte count. In our assay, the derived allele at rs56094005 (Anc/Der; C/G; haplotype frequency = 8.7% in EstBB) reduced activity (**Fig. 6d**) compared to the ancestral haplotype (91% frequency in EstBB; **Fig. 6e**). This was consistent with its eQTL direction of downregulating *GFRA2*. While the double derived haplotype maintained similar activity to ancestral having the rare, derived allele at rs373143737 alone (Der/Anc) increased activity. Only rs373143737 overlaps a predicted ATMIN motif (**Fig. 6f**), while the 57 bp between contains 9 additional motifs. This includes motifs such as SP3, KLF6, KLF7, MAZ, and several other zinc-finger motifs (**Fig. 6f**).

A contrasting example involved two variants only 12 bp apart within a broadly active enhancer. rs11694743 (chr2:127842607, A>G) and rs549250007 (chr2:127842619, G>A), separated by only 12 bp. In our MPRA, the ancestral haplotype and the haplotype carrying only the derived allele of rs11694743 (Der/Anc) showed similar regulatory activity. In contrast, introduction of the derived allele of rs549250007 reduced activity. With the strongest reduction observed when both derived alleles were present, indicating that the effect of rs549250007 depended on the allelic background at rs11694743 (**Fig. 6g**). In EstBB, the ancestral and Der/Anc haplotypes were common, with frequencies of 38.6% and 60.5%, respectively, whereas haplotypes carrying the derived allele of rs549250007 were rare. Despite their close proximity, the two derived alleles did not commonly occur together in EstBB, consistent with the very low R² between the variants (R² = 0.012; D′ = 0.925) (**Fig. 6h**). rs11694743 is also an eQTL whose derived G allele is associated with increased *BIN1* expression, although this allele alone had little effect on regulatory activity in our MPRA. Both variants occur within a dense motif context and together overlap 13 unique predicted TF motifs. Four motifs, ZNF746, ZNF48, MBD3, and KMT2A, overlap both variant positions, while additional motifs overlap only one of the two variants (**Fig. 6i**). Overall, the activity pattern suggests that rs549250007 is the primary driver of reduced regulatory activity, while the derived allele at rs11694743 further modifies and strengthens this effect in the double-derived haplotype. Together, these examples illustrate how non-additive regulatory effects can arise in different local sequence and population haplotype contexts.

## Discussion

Understanding how non-coding genetic variation contributes to complex traits and disease remains a major challenge. GWAS and eQTL studies have identified thousands of associations but often cannot identify the causal variants or explain the regulatory mechanisms underlying these associations. Large-scale functional assays such as MPRAs have therefore become an important approach for experimentally testing variant function and complementing statistical and computational methods. To date, most MPRA studies have focused on testing variants individually and assessing the effects of each allele in isolation (Capauto et al., 2024; Griesemer et al., 2021; Jiang et al., 2024; Liu et al., 2017; Tewhey et al., 2016; Ulirsch et al., 2016). Here, we instead tested all four haplotype combinations of 7,285 pairs of closely spaced variants identified from EstBB WGS data. This haplotype-based design captures a layer of regulatory complexity that single-variant assays cannot directly measure. In K562 cells, 57% of tested haplotype sets contained at least one DA haplotype. Because our library was enriched for variants with regulatory potential in blood-related enhancers, this proportion should not be interpreted as representative of regulatory variants genome-wide. The regulatory effects associated with individual variants often depended on the neighboring allelic background. Variants could retain, lose, or in some cases reverse their effects when paired with a different neighboring allele. These findings support previous observations that the functional effect of a regulatory variant cannot always be inferred from the variant alone, because its surrounding sequence and genetic context can modify its effect (Chandler et al., 2017; Maricque et al., 2017; Sackton & Hartl, 2016). Recent endogenous editing studies similarly demonstrate that the effects of regulatory sequence changes can be strongly dependent on local genomic and cellular context (Martyn et al., 2025).

Testing all four haplotypes also allowed us to determine whether the effects of nearby variants combined additively. Within our HC subset, 59% (105/178) of variant pairs showed non-additive effects. This proportion should not be interpreted as an estimate of the genome-wide prevalence of regulatory non-additivity, because the HC subset was deliberately enriched for haplotype sets with substantial regulatory activity and differential effects. Nevertheless, our results add to growing evidence that multiple nearby functional variants can contribute to the same regulatory locus. Abell et al. (Abell et al., 2022) showed that 17.7% of tested eQTL loci contained more than one variant with an allelic effect in tight LD. In an analysis of 2,097 pairs separated by less than 75 bp, they found that most haplotype effects were additive but that non-additive interactions also occurred between the two variants. More recently, Siraj et al. (Siraj et al., 2026) tested all four haplotypes of nearby fine-mapped regulatory variant pairs within the same CRE and showed non-additive regulatory effects in 11% of tested pairs. Notably, they also showed that in non-additive pairs variants were located closer together than additive pairs. In majority of those pairs combinations of two activity-increasing alleles commonly had smaller effects than expected from their individual effects. We observed both patterns independently; non-additive pairs were closer than additive variant pairs, and double-derived haplotypes often had smaller effects than expected from the sum of the two individual effects. Interestingly, variant distance was not associated with the likelihood or magnitude of DA across the library. These findings suggest that proximity may be more important for how two regulatory variants interact than for whether an individual variant has a detectable regulatory effect. The predominantly sub-additive effects could reflect nonlinear or saturating regulatory responses, in which one sequence change already produces much of the achievable change in activity and the second therefore contributes less than expected. For pairs with effects in opposing directions, partial compensation may also resemble linkage masking, whereby variants occurring on the same haplotype reduce one another’s phenotypic or regulatory effects (Siraj et al., 2026; Zhang et al., 2026).

We also examined population-level characteristics that could explain the regulatory effects observed in our library. Rare haplotypes (frequency <1% in EstBB WGS data) were associated with a modest increase in the probability of DA, while effect magnitude was not associated with frequency. A similar pattern was observed at the level of individual allele frequency, with rarer alleles more frequently associated with differential activity. This is consistent with previous work showing that rare variants are enriched for functional effects (Zhu et al., 2011), and that rare regulatory variants can alter gene expression (Li et al., 2014, 2017; Montgomery et al., 2011; Zhao et al., 2016). Recent population-scale studies further demonstrate the importance of population-specific rare non-coding variation for quantitative traits and show that such variants can have substantial functional effects (Koyama et al., 2024). Importantly, however, neither allele nor haplotype frequency was associated with the magnitude or direction of DA among haplotypes that were already DA. Thus, in our data, rarity predicts the likelihood of a detectable regulatory effect rather than the size of that effect once present. The modest enrichment of DA among rare haplotypes is consistent with the possibility that purifying selection contributes to keeping regulatory variants with functional consequences at lower frequencies (Glassberg et al., 2019). However, our data does not directly demonstrate selection. Similarly, among the HC non-additive sets, haplotypes not observed in EstBB were frequently DA and often double-derived. Furthermore, double-derived DA haplotypes were, on average, negative, indicating a possible disruption of the regulatory function of the ancestral haplotype. Because haplotype frequencies were estimated from 2,420 WGS samples, haplotypes not observed in this dataset may still occur at very low frequencies in the broader population. Their low frequency could also reflect recent origin, population history, or constraints imposed by LD and recombination. Therefore, distinguishing these possibilities from purifying selection will require additional population-genetic analyses and larger population samples.

TF binding provides one possible mechanism through which nearby regulatory variants shape regulatory function. Our motif analysis showed that additive and non-additive sets had similar overall motif density and were similarly likely to contain at least one variant overlapping a predicted TF motif. However, the organization and combination of these overlaps differed. Additive sets had more unique motifs overlapping one of the assayed variants. They also had a more diverse set of motif combinations, including cases in which the two variants overlapped distinct motifs. Non-additive sets, in contrast, had more often a motif overlapping only one variant or a motif shared by both variants. We did not observe non-additive sets in which both variants independently overlapped distinct motifs. These results suggest that combination of motifs rather than overall motif density, may contribute to whether the effects of nearby variants combine additively. This interpretation is consistent with experimental studies showing that TF motif activity is strongly influenced by surrounding sequence, neighboring motifs, and their spatial organization (Morgunova & Taipale, 2017; Rao et al., 2021; Reiter et al., 2023). Other work shows that combinations of TFs can establish cooperative regulatory environments in which the effect of one binding site depends on other factors present in the enhancer (Kribelbauer-Swietek et al., 2024). At the same time, motif overlap alone should not be interpreted as evidence of TF binding or as proof of the molecular mechanism. Siraj et al. (Siraj et al., 2026) for example, found that only 69% of their high-confidence regulatory variants could initially be explained by disruption of a known TF motif. Saturation mutagenesis then showed additional sequence features and candidate TFs for many variants lacking a motif-based explanation. This is particularly relevant to our chromosome 8 example, in which rs56094005 altered regulatory activity despite not directly overlapping a predicted TF motif. This suggests that regulatory effects can arise through changes in broader sequence context or through binding sites not captured by current motif predictions.

Several of the non-additive sets in our study were in a locus that is associated with human traits, illustrating how these interactions may complicate interpretation of GWAS and eQTL signals. Thirteen of the 105 HC non-additive sets contained at least one GWAS-associated variant. One example involved two adjacent variants on chromosome 17 that overlapped the same predicted ZNF184 motif. Introducing either derived allele reduced regulatory activity, while combining the two produced little additional effect. This provides a simple example of how two variants affecting the same local regulatory feature could result in sub-additive response. Both derived alleles are associated with increased IKZF3 expression in eQTLGen dataset, whereas they reduced regulatory activity in our MPRA. IKZF3 encodes a TF with an important role in lymphocyte development and B-cell differentiation (Morgan et al., 1997; Wang et al., 1998). It has also been a target for multiple myeloma (MM) therapies, where elevated *IKZF3* expression acts like pro-survival factor, reducing the effectiveness of the treatment (Chowdhury et al., 2024).

Rather than providing a direct mechanistic link between MPRA activity and *IKZF3* expression, the difference in direction between the MPRA and eQTL effect highlights the importance of endogenous context. Our MPRA measures the regulatory potential of a 170-bp sequence in a reporter system, and sequence length itself can substantially influence MPRA measurements (Klein et al., 2020). Regulatory effects in the endogenous genome may additionally depend on surrounding sequence features, chromatin state, enhancer–promoter interactions, and cellular context. Endogenous regulatory editing studies similarly show that the effects of regulatory variants can depend on both genomic and cellular context (Martyn et al., 2025).

The other trait-associated examples further showed that non-additive regulatory effects can occur across different sequences and LD contexts. At chromosome 8, the two variants were separated by 57 bp and only one directly overlapped a predicted TF motif. At the same time, variants on chromosome 2 were only 12 bp apart, and within this 12 bp was a dense cluster of overlapping motifs. These two examples had very different population structures. Variant pairs on chromosome 2 showed very low correlation despite their close proximity, illustrating that physical distance alone does not determine LD or the haplotypes in which regulatory variants occur. Together, these examples suggest that non-additive regulatory effects can emerge from different combinations of local sequence architecture and population haplotype structure.

Here we show the importance of moving beyond single variant testing when interpreting the regulatory effects of non-coding variation. By testing all haplotype combinations of close proximity variant pairs we gain better understanding how non-coding variants together affect gene regulation as we saw individual variant effects often depended on the neighboring allelic background. Among haplotype sets with strong regulatory effects, combined effects often deviated from additivity. The association of non-additivity with variant proximity and differences in motif architecture suggests that local sequence organization contributes to these interactions. The population analysis shows that rare alleles and haplotypes are more likely to have regulatory effects. Together, our results demonstrate that the functional consequences of regulatory variation can depend on the haplotypes in which variants occur. This supports haplotype-based functional assays as an important complement to single-variant approaches for interpreting non-coding variants. Future studies should combine such assays with endogenous genome editing, direct measurements of TF binding and use larger population datasets. This will help to determine how these interactions combine in their native genomic and population context.

## Acknowledgements

This work has received funding from European Union’s Horizon 2020 research and innovation programme under grant agreement No 810645 and by the European Union through the European Regional Development Fund (grant no MOBEC008). This work has also been supported by Estonian Research council grant PRG3113. The research was conducted using the Estonian Center of Genomics/Roadmap II funded by the Estonian Research Council (project number TT17). Data analysis was carried out in part in the High-Performance Computing Center of University of Tartu. St Vincent’s Institute acknowledges the infrastructure support it receives from the National Health and Medical Research Council Independent Research Institutes Infrastructure Support Program and from the Victorian Government through its Operational Infrastructure Support Program.

## Materials and methods

### MPRA design and library preparation

#### MPRA variant selection

We designed the MPRA library to evaluate the regulatory effects of close-proximity variant pairs located in enhancers. We selected variants from whole-genome sequencing (WGS) data of 2,420 individuals from Estonian Biobank (EstBB) (Leitsalu et al., 2015; Milani et al., 2025). We kept SNVs with a minor allele count (MAC) >3 and genotype missingness rate of <10%. Filtered SNVs were overlapped with ChromHMM enhancer annotations (states 6_EnhG and 7_Enh) from the Roadmap Epigenomics Project (Bernstein et al., 2010; Roadmap Epigenomics Consortium et al., 2015). We had two enhancer-based strategies: i) blood specific enhancers, variants in sequences that were annotated as enhancers in at least 2 blood cell types, with at least one of them having to be a B-cell (E031 or E032) and had no annotation as enhancers in any other non-hematopoietic cell types; and ii) broadly active enhancers, that were annotated as enhancers in more than 85 Roadmap cell types. For both of those selections, additional filtering was applied, the 1000 Genomes Project (Auton et al., 2015) strict callability mask. Variants that were common in Estonian population (MAF > 10%) but not present in 1000 Genomes Phase 3 dataset were excluded. The SNV pairs passing the filtering and separated by up to 75 bp, included at least one cis-eQTL (-log10(p) ≥ 15) identified in eQTLGen v1(Võsa et al., 2021). We annotated the selected SNVs as “ancestral” and “derived” based on the reconstructed ancestral genome sequences from Ensemble database (release 115) (Dyer et al., 2025; Paten et al., 2008). After applying the set of these criteria, we had 12,686 SNVs within blood specific enhancers and 1,884 SNVs within enhancers mapped in >85 Roadmap tissues. A set of negative (n = 212) and positive controls (n = 219) for this assay were selected from a previous study (Tewhey et al., 2016). Additionally, we selected 129 expression positive variants from the same study and scrambled their sequences, to use them as additional set of negative controls. iPSC (n = 150; Roadmap Epigenomics data codes: E020, E019, E018, E021, E022) and brain (n = 150; Roadmap Epigenomics data codes: E071, E074, E068, E069, E072, E067, E073, E070, E082, E081) cell line specific enhancers selected from the Roadmap Epigenomics data (Bernstein et al., 2010; Roadmap Epigenomics Consortium et al., 2015). In total our library consisted of 30,000 unique sequences (Table S1).

#### MPRA oligonucleotide library design and synthesis

Oligos were synthesized by Twist Bioscience as 200 bp sequences containing 170 bp of genomic sequence flanked by 15 bp adapter sequences on each end (5’ACTGGCCGCTTGACG [170 bp oligo] CACTGCGGCTCCTGC3’). The oligo pool was dissolved in TE buffer in final concentration 20 ng/µl according to the Twist Bioscience DNA resuspension guidelines.

A round of PCR was carried out to amplify and convert the oligos into double stranded DNA using primers complementary to the adapter sequences (F_adapter and R_adapter; Table S7). We amplified the library following the Twist Bioscience PCR amplification protocol, using KAPA HiFi HotStart Polymerase (KK2502, Roche) with 12 cycles in 6 parallel reactions. Samples were pooled and purified with QIAquick PCR Purification Kit (28104, Qiagen) according to the manufacturer’s protocol. We quantified DNA concentrations with NanoDrop/QBit (Thermo Scientific). Fragment size we assessed with TapeStation4200 (G2991BA, Agilent) to confirm that the PCR product corresponds to the correct fragment length after amplification.

A second round of PCR was performed to add 15 bp unique barcodes, HiFi assembly overhangs, *BsiWI-HF* and *SfiI* restriction sites using F_PCR and barcoding_R primers (Table S7). Eight parallel reactions were carried out with Q5 High-Fidelity 2X Master Mix (M0492L, NEB). Samples were pooled and purified with QIAquick PCR Purification Kit (28104, Qiagen) according to manufacturer’s protocol. Concentrations were quantified with NanoDrop/QBit (Thermo Scientific). Fragment size was assessed using TapeStation4200 (G2991BA, Agilent) to confirm that the PCR product corresponds to the correct fragment length after amplification.

#### MPRA vector assembly

Barcoded oligos were digested with *SfiI* (R0123L, NEB) according to manufacturer’s protocol and purified with QIAquick PCR Purification Kit (28104, Qiagen). Concentration was quantified with NanoDrop/Qbit (Thermo Scientific), and fragment size distribution was verified with TapeStation4200 (G2991BA, Agilent). The pMPRA1 (49349, Addgene) plasmid was similarly digested with *SfiI* (R0123L, NEB) and purified with 0.5x AMPure XP beads (A63881, Beckman Coulter).

Barcoded oligos were ligated into *SfiI* (R0123L, NEB) digested pMPRA1 backbone using T4 DNA ligase (M0202M, NEB) at 1:3 plasmid insert ratio, with incubation at 16 °C overnight. Ligated products were transformed into Promega JM109 competent cells (L2005, Promega) by heat shock. Following recovery in SOC medium at 37 °C for 1h, cells were plated on LB agar plates with serial dilutions. Library complexity was estimated >10^8^ CFUs. To validate successful ligation, ten colonies from dilution plates were selected and check by colony PCR to confirm the insertion of oligos. Colonies were collected from the LB plates and plasmid was purified using Zymo Pure II Plasmid Maxiprep kit (D4203, Zymo Research), eluting twice with 400 µl of EB buffer (total 800 µl).

To associate unique barcodes to their corresponding oligos, 10 ng of pMPRA1 plasmid library containing barcoded oligos (pMPRA1:oligo) was PCR amplified (12 cycles) with KAPA HiFi HotStart Polymerase (KK2502, Roche) using F_adapter and buffer_R (Table S7). Products were sequenced on a NovaSeq X Plus (Novogene) to a depth of 300 million 150 bp paired-end reads. Barcode-oligo matching was performed using MPRAmatch pipeline by Tewhey et al. (Tewhey et al., 2016). After sequencing, 29,484 (98.3%) of the initial 30,000 sequences were detected, of which 843 were controls and 28,641 were the haplotype sequences of selected variant pairs.

For the final construction, 20 ug of the pMPRA1 plasmid library containing barcoded oligos (pMPRA1:oligo) was digested with *BsiWI-HF* (R3553L, NEB), purified with GeneJet PCR Purification Kit (K0702, Thermo Scientific) and eluted in 50 µl of EB. A synthetic gBlock (IDT) containing a minimal promoter (minP), eGFP open reading frame and cloning overhangs (Table S7) was inserted into *BsiWI-HF* digested pMPRA1:oligo library using the NEBuilder HiFi DNA Assembly at a1:3 ratio (M0541L, NEB). The reaction was incubated at 50 °C for 90 min, followed by 1:1 bead clean up (A63881, Agencourt) and eluted in 26 ul of EB. This was the final library, where the pMPRA1 plasmid contained the designed oligos with minimal promoter and eGFP open reading frame (pMPRA1:oligo:egfp).

To generate transfection-ready MPRA libraries, 10 μL of pMPRA1:oligo:egfp plasmid was electroporated (2 kV, 200 ohm, 25 μF) into 100 μL of 10-beta electrocompetent *E. coli* cells (C3020K, NEB). Electroporated cells were divided across 5 tubes, each recovered in 1 ml of SOC medium for 1 hour at 37°C, and expanded in 500 ml of LB (5 x 500 ml, 2 ml per 500 ml) supplemented with 100 μg/ml of ampicillin overnight at 37 °C. For each aliquot, serial dilution was plated following SOC recovery to estimate library complexity (>10^8^ CFUs). Ten colonies were again screened by colony PCR to verify eGFP insertion. For plasmid purification, cells were centrifuged and wet pellet was weighted, plasmid was purified using Qiagen Plasmid Giga Kit (12192, Qiagen). Plasmid extractions were pooled and normalised to 1 ug/ul.

### Cell transfection and data generation

#### Transfection of MPRA constructs into target cells

Lymphoblastoid cells (GM12878, Coriell) and myelogenous leukemia cells (K562) were maintained in RPMI 1640 (61870, Life Technologies) supplemented with 15% FBS (A5670701, Gibco) at a cell density of 2-10×10^5^ cells/ml. 24h prior to transfection, cultures were split and supplemented with fresh cell culture media.

On the day of transfection, cells were counted and collected by centrifugation at 120 x g for 5 minutes, then resuspended in Neon NxT buffer R. Electroporation was performed in 100 μl volumes using the Neon NxT Electroporation system (N10096, Life Technologies). For GM12878, 3 pulses of 1300V 10ms each were applied, while for K562 3 pulses of 1450V 10ms each were used. In each cell line, 80 x 10^6^ cells were transfected with 400 ug of pMPRA1 library. Cells were recovered in 100 ml of RPMI supplemented with 15% of FBS for 24h at 37°C.

Following recovery cells were collected by centrifugation (120 x g, 5 minutes), washed with 2 mL DPBS (141900144, Thermo Fisher Scientific), and pelleted again. 90% of the cells were used for RNA extraction and two aliquots of 5% were used for DNA extraction. Cells were collected again with centrifugation 120 x g for 5 minutes, DPBS was removed, and the cell pellet was stored at –80 °C. For both cell lines, 7 replicates were performed.

#### RNA extraction and cDNA synthesis

Total RNA was extracted using the RNeasy Maxi Kit (75162, Qiagen), with the on-column DNase digestion according to the manufacturer’s protocol. A second DNase treatment was performed on purified RNA using TURBO DNA-free Kit (AM1097, Thermo Fisher Scientific), and the reaction was stopped by adding 0.1 volume of DNase inactivation reagent. DNA was extracted by using DNeasy Blood & Tissue Kit (69504, Qiagen). First-strand cDNA was synthesized from the DNase treated RNA using SuperScript IV (18090050, Invitrogen) and a primer specific to the 3’ UTR (Buffer_R, Table S7). To determine the optimal number of PCR cycles for library amplification, qPCR was performed on both pDNA and cDNA samples. Reactions were carried out in triplicates with HOT FIREPol EvaGreen qPCR Supermix (08-36-00001, Solis BioDyne) with primers complementary to buffer sequences and GFP (RNAseq_F and Buffer_R; Table S7).

The cDNA and pDNA libraries were amplified by using the ct values determined by qPCR with Q5 High-Fidelity DNA Polymerase (M0491L, NEB; RNAseq_F and buffer_R; Table S7). PCR products were purified using Monarch PCR & DNA Cleanup Kit (T1130L, NEB) and validated with TapeStation4200 (G2991BA, Agilent). Concentrations were quantified using NanoDrop/QBit (Thermo Scientific).

To confirm library quality, initial sequencing of cDNA and pDNA was performed at a depth of 2 x 20 million 150 bp paired-end reads (NovaSeq X Plus, Novogene). After verifying sample quality, additional cDNA and pDNA sequencing was performed to a depth of 2 x120 million 150 bp paired-end reads.

### MPRA data processing and main analysis

Oligo-barcode association was performed following the MPRAmatch pipeline from Tewhey et al., 2016. A custom Python script by Navya Shukla was used in place of pull_barcodes.pl to match our library design. Barcode counts per unique oligo were obtained using MPRAcount workflow by Tewehy et al., 2016. Paired-end 150 bp reads were merged with Flash v2.2 (Magoč & Salzberg, 2011) and reads from the two rounds of sequencing were merged. The oligo-barcode dictionary from the previous step was used to map the reads and generate count table for cDNA and pDNA libraries **(Table S2).** This table contain unique barcodes per each oligo and the counts per barcodes and was use for downstream analysis performed in R (version 4.5.1).

Barcode counts per oligo were aggregated and to identify haplotypes with regulatory activity and DA, we followed the pipeline described by Abell et al. (Abell et al., 2022) using the R package DESeq2 (Love et al., 2014) as the framework for normalization and statistical testing. For both libraries, counts were normalized across replicates using DESeq2’s median ratios method. For activity analysis and DA analysis, only oligos with a mean raw count of >400 across all samples were retained. In addition, haplotype sets that did not contain read counts for all 4 haplotypes were removed from the analysis. For K562 we retained 5,988 full haplotype sets for further analysis and for GM12878 6,106 haplotype sets. In total, combined over the two cell lines 6,168 haplotype sets were analyzed for activity and DA.

To assess significant differences between haplotypes in DNA and RNA counts, a negative binominal generalized linear model (GLM) was fitted within DESeq2, with independent dispersion estimates for each cell type and library. To estimate haplotypes activity, RNA counts were compared to DNA counts, by including contrast in the design matrix (RNA vs DNA). The following design formula was used for activity analysis: ∼material + replicate. For differential activity analysis, within each haplotype set (4 haplotypes: Anc/Anc, Anc/Der, Der/Anc and Der/Der), each non-reference haplotype activity was compared against ancestral haplotype (Anc/Anc) in that group for DA by including the contrasts in the design matrix of DESeq2. For that we used the following design formula: ∼material + ancestry_combination + material:replicate + material:ancestry combination. We assessed statistical significance using Wald’s test, and p-values were adjusted for multiple testing using Benjamin-Hochberg’s false discovery rate (FDR) procedure. We considered haplotypes with an adjusted p-value <0.01 differentially active relative to the ancestral (Anc/Anc), reference haplotype (Table S3, S4). In addition, we generated a high-confident (HC) DA haplotype set for which we applied additional filters. For this set we required at least 1 haplotype out of 4 to show regulatory activity, defined as an RNA to DNA ratio of |log_2_FC RNA/DNA >1| in the same group. Second, at least one of the three non-ancestral haplotypes (Anc/Der, Der/Anc, Der/Der) was required to show absolute DA of >0.5. Additionally, the DA had to be statistically significant, with an adjusted p-value of <0.01 (Table S5).

To test non-additive interactions between variants, we compared the observed DA of the double derived haplotype (Der/Der) to the expected additive DA derived from the sum of DA of the two single derived haplotypes (Anc/Der, Der/Anc). Uncertainty in the additive expectation was estimated by combining the standard errors of the two single derived haplotypes. We then calculated 95% confidence intervals for both the observed and expected DA (Table S5). We classified haplotypes as non-additive when these confidence intervals did not overlap and as additive when they did.

#### Genomic annotation and variant to gene mapping

Genomic annotation of tested variants was per formed using the R (version 4.5.1) packages rtracklayer and GenomicRanges (Lawrence et al., 2013). Transcription start site (TSS) were defined in a strand-specific manner using GENCODE v48lift37 gene models (GTF file: gencode.v48lift37.annotation.gtf.gz, GRCh37/hg19). For genes on the positive strand, the TSS was defined as the gene start coordinate. For the genes on the negative strand, the TSS was defined as the gene end coordinate, matching the 5’ end of each gene model. For each MPRA variant pair, the midpoint between the two variant positions was calculated and the distance to the nearest annotated TSS was identified using the nearest() function from GenomicRanges. Signed distance to the nearest TSS were calculated relative to gene strand orientation and shown in kilobases. Variants were then binned by distance to the nearest TSS. We also obtained hg38 genomic coordinates for our variants using the liftOver function from rtracklayer package (Lawrence et al., 2009) in R. For the liftover we used hg19ToHg38.over.chain file from UCSC Genome Browser (Casper et al., 2026).

#### Overlap with open chromatin

HC haplotype set overlap with open chromatin regions were assessed using ATAC-seq and DNase I hypersensitivity (DHS) data from K562 cells. ATAC-seq peaks (ENCODE accession ENCSR483RKN) and DHS peaks (ENCODE accession ENCSR558GSX) were imported and converted into genomic ranges using the GenomicRanges package (Lawrence et al., 2013). Since ATAC-seq and DHS files were in hg38, we used the hg38 co-ordinates for overlaps. For each DA haplotype we took the midpoint between the two variants and overlaps with ATAC-seq and DHS peaks were identified using genomic interval intersection.

#### Population specific haplotype analysis

Because the MPRA constructs were designed to include all possible allele combinations for the variant pairs, we used PLINK2 (Chang et al., 2015) to estimate haplotype frequencies for the Estonian population using whole-genome sequencing (WGS) data (n = 2,420) from the Estonian Biobank (EstBB) (Leitsalu et al., 2015; Milani et al., 2025). The variants tested in this study were extracted from EstBB WGS dataset using bcftools/1.21 (Danecek et al., 2021). Extracted variants were converted into PLINK2 format, and for each variant pair linkage-disequilibrium (LD) measures and haplotype frequencies were calculated.

#### Transcription factor binding motif and occupancy

In order to find the best match for a human transcription factor (TF) motif, sequences were analyzed using the FIMO tool (Grant et al., 2011) from MEME suite. The HOCOMOCO v14core (Vorontsov et al., 2024) of human TF binding motifs was used as the reference motif set. We were looking for the best match for a human TF motif in HOCOMOCO. For each DA haplotype sequence identified in the MPRA, the corresponding genomic sequences were scanned for TF motif occurrences using FIMO with default parameters, with the p-value threshold of <0.0001 to identify significant matches. To assess allele-specific motif disruption or gain, both reference and alternative allele haplotypes were analyzed. The output from FIMO was then overlapped with the variant positions to accurately find motifs that overlap with the tested variants. To find matches, (i) we limited the list to motifs with model “0”; (ii) TF motif classes “A” and “B”; (iii) applied q-score threshold of <0.05.

#### GWAS association

To determine how many of our tested variants in our non-additive haplotype sets have a known trait association, we used the GWAS Catalog (Cerezo et al., 2025; Sollis et al., 2023). From the GWAS Catalog we kept only associations with the genome-wide significance threshold (p < 5 x 10^-8^). We then overlapped variants in our non-additive set with this filtered list by genomic position (GRCh38/hg38).

