## Supplementary figures S1-S9 for "Massively parallel characterization reveals context-dependent and non-additive regulatory effects of closely spaced variant pairs"

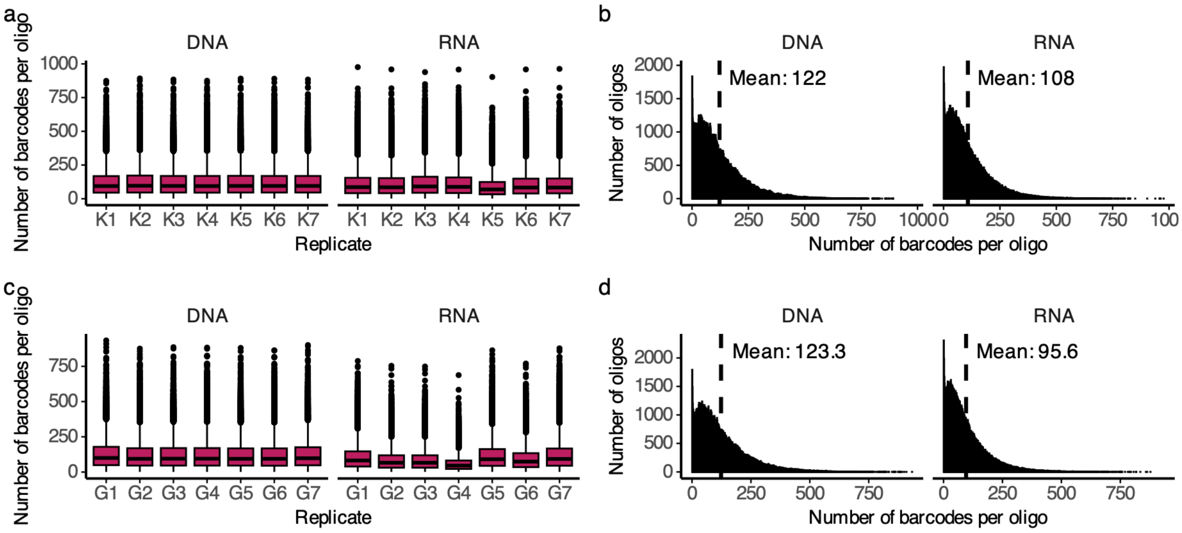


**Supplementary Fig. S1 Barcodes per oligo across K562 and GM12878 replicate libraries. a** Barcode counts across K562 DNA and RNA libraries. **b** Number of average barcodes per oligos in K562 DNA and RNA libraries. **c** Barcode counts across GM12878 DNA and RNA libraries. **d** Number of average barcodes per oligo in GM12878 DNA and RNA libraries.

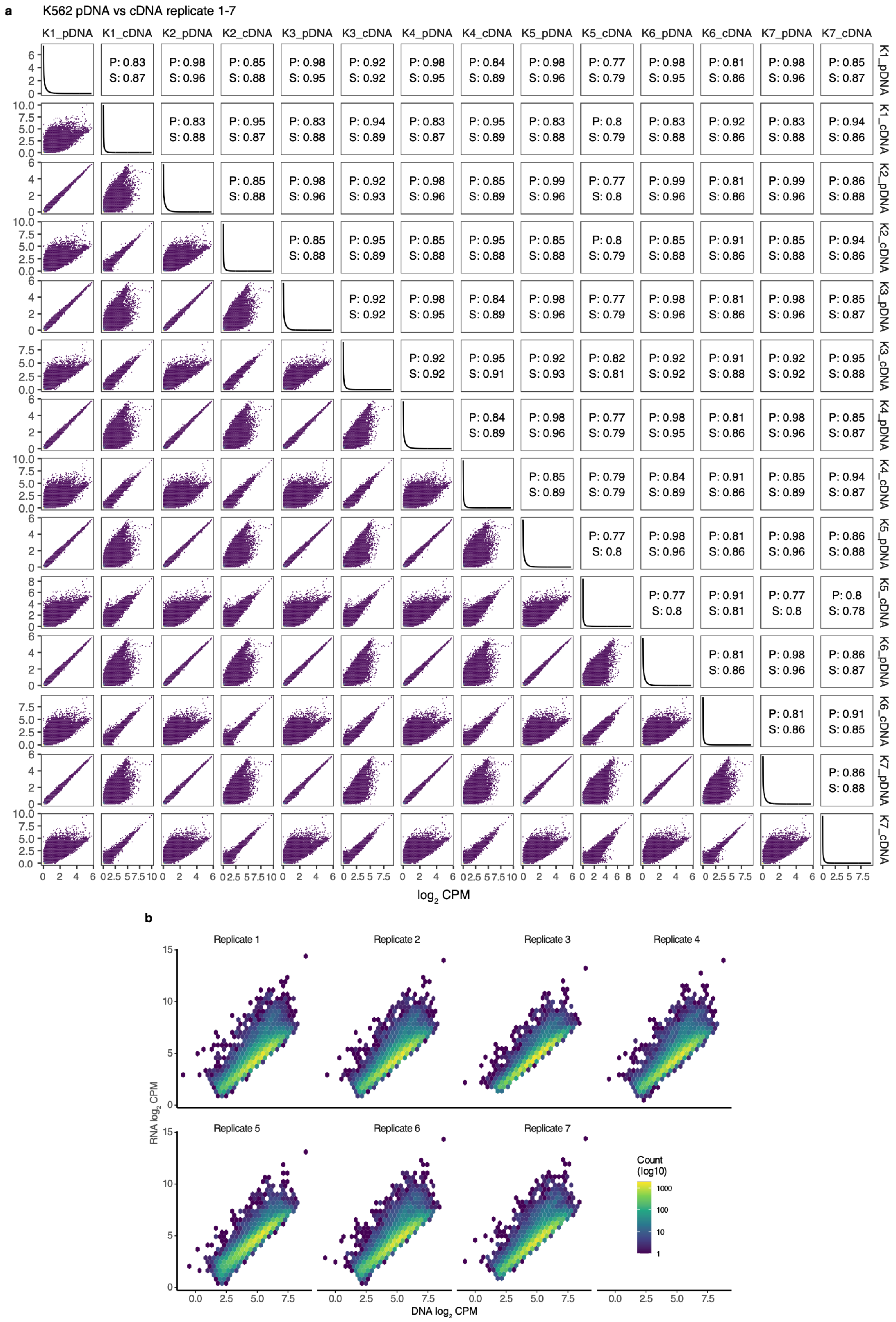


**Supplementary Fig. S2 Correlation between cDNA and pDNA replicates in K562. a** full correlation matrix between K562

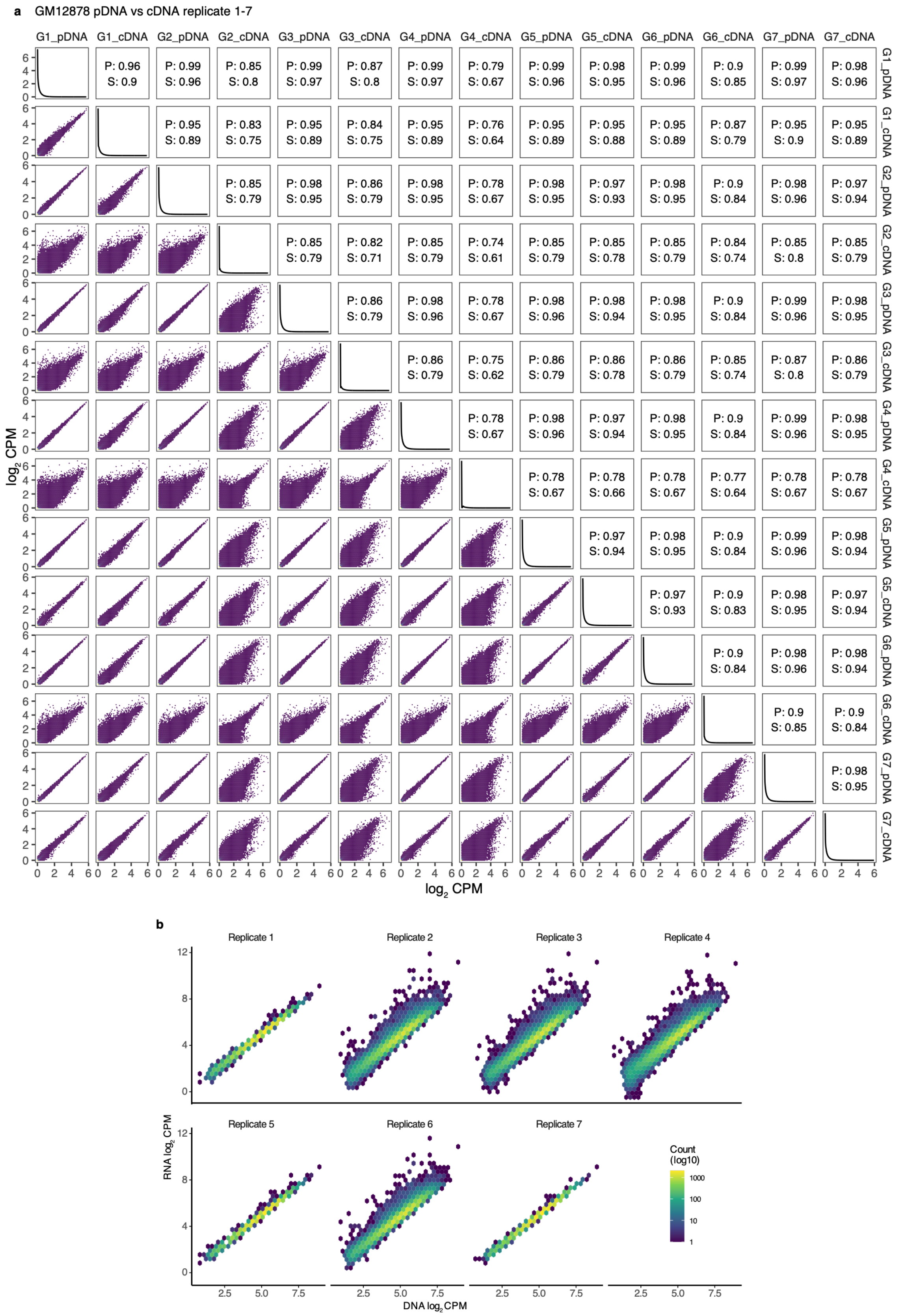


.

**Supplementary Fig. S3 Correlation between cDNA and pDNA replicates in GM12878. a** Full correlation matrix between GM12878 pDNA and cDNA replicates. **b** Replicate specific correlation of cDNA and pDNA.

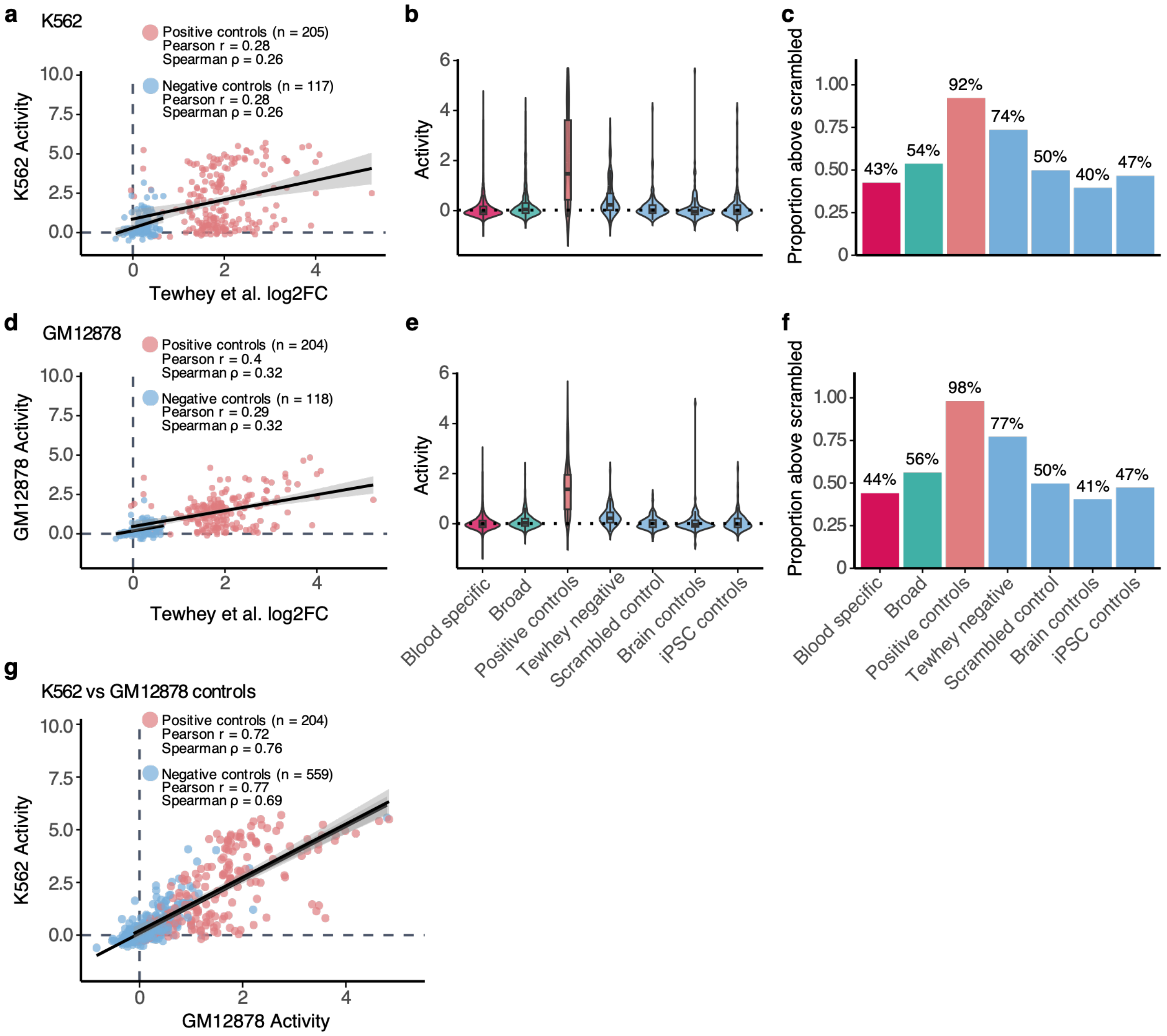


**Supplementary Fig. S4 Activity ($\log_2$FC RNA/DNA) distribution across cell lines and sets. a** Correlation of positive and negative control activity between original Tewhey et al. results and our K562 MPRA results. Correlation was calculated separately between positive controls and negative controls. **b** Activity in K562 with positive and negative controls, with dotted line indicates scrambled control median activity. **c** Proportion of haplotype sets and controls with DA and activity levels above scrambled control median activity. **d** Correlation of positive and negative control activity between original Tewhey et al results and our GM12878 MPRA results. **e** Activity in GM12878 with positive and negative controls, with dotted line indicates scrambled control median activity. **f** Proportion of haplotype sets and controls with DA and activity levels above scrambled control median activity. **g** Correlation of positive and negative controls between GM12878 and K562 in our experiment. Correlations were calculated separately for both groups.

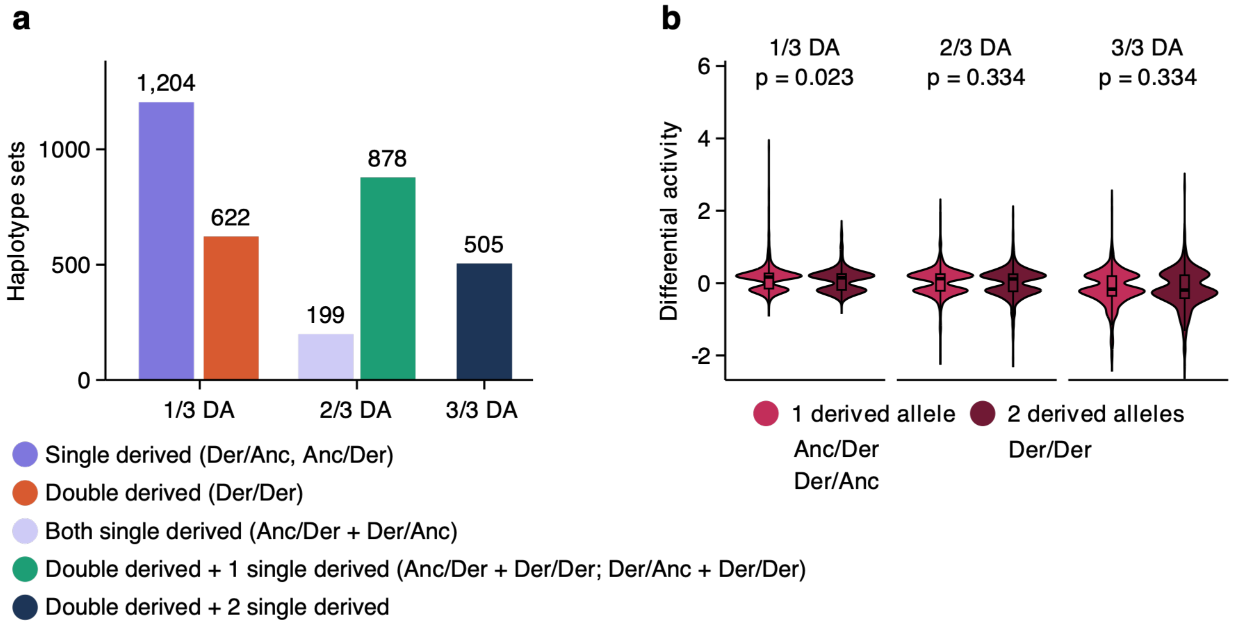


**Supplementary Fig. S5 DA haplotype set composition and DA by derived allele dosage. a** Number of haplotype sets in DA category split by DA haplotype combinations. **b** DA stratified by number of derived alleles in DA haplotypes. For each DA category (1/3, 2/3, 3/3) Anc/Der and Der/Anc haplotypes were categories as “1 derived allele” and Der/Der haplotypes as “2 derived alleles”.

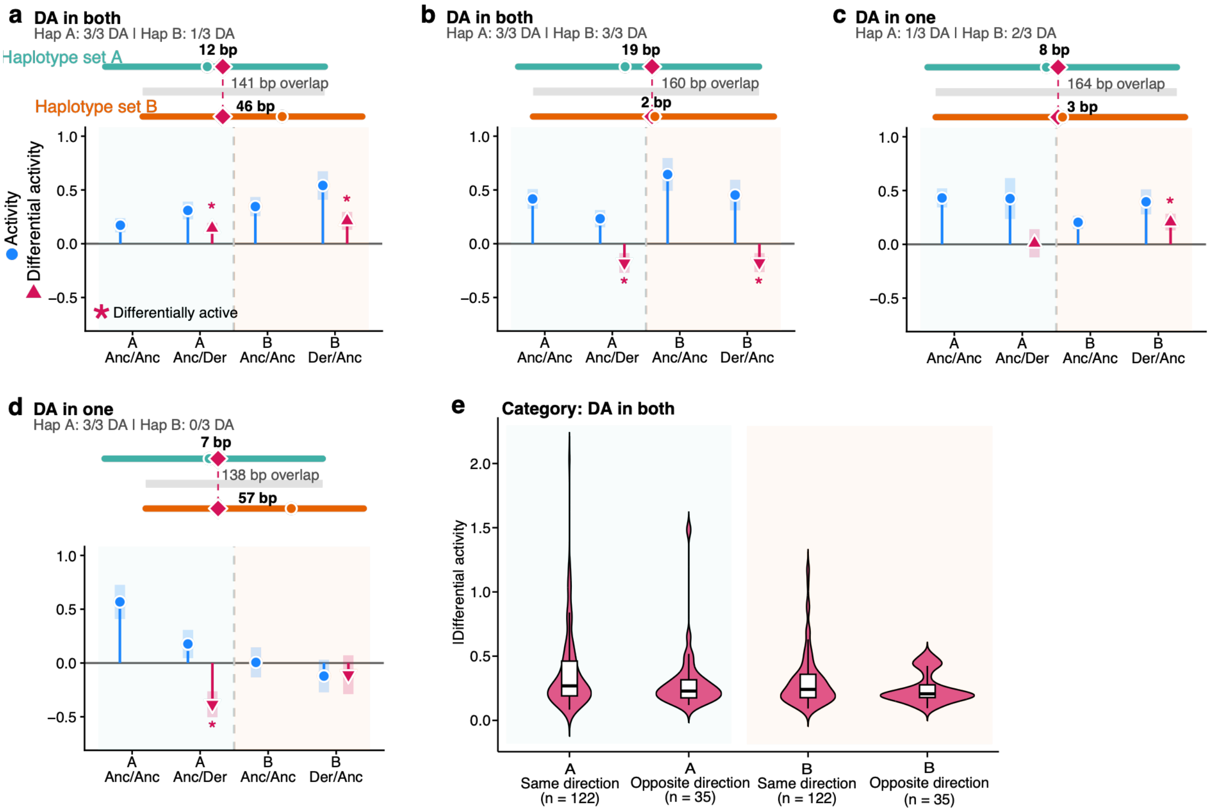


**Supplementary Fig. S6. Examples of the variants tested across multiple haplotypes sets with new variant pair. a-b** 2 examples from the "DA in both" category, where in both haplotypes variant increase expression compared to ancestral haplotype (a); variant decreases expression (b) and variant is DA in both but has opposite DA direction (c). c-**d** are 2 examples from "DA in one" category. e DA in the "DA both" category. Shown is the magnitude of DA for the variants that maintained the same DA direction in both backgrounds and for variants that were DA in both tested haplotype sets but had opposite DA direction.

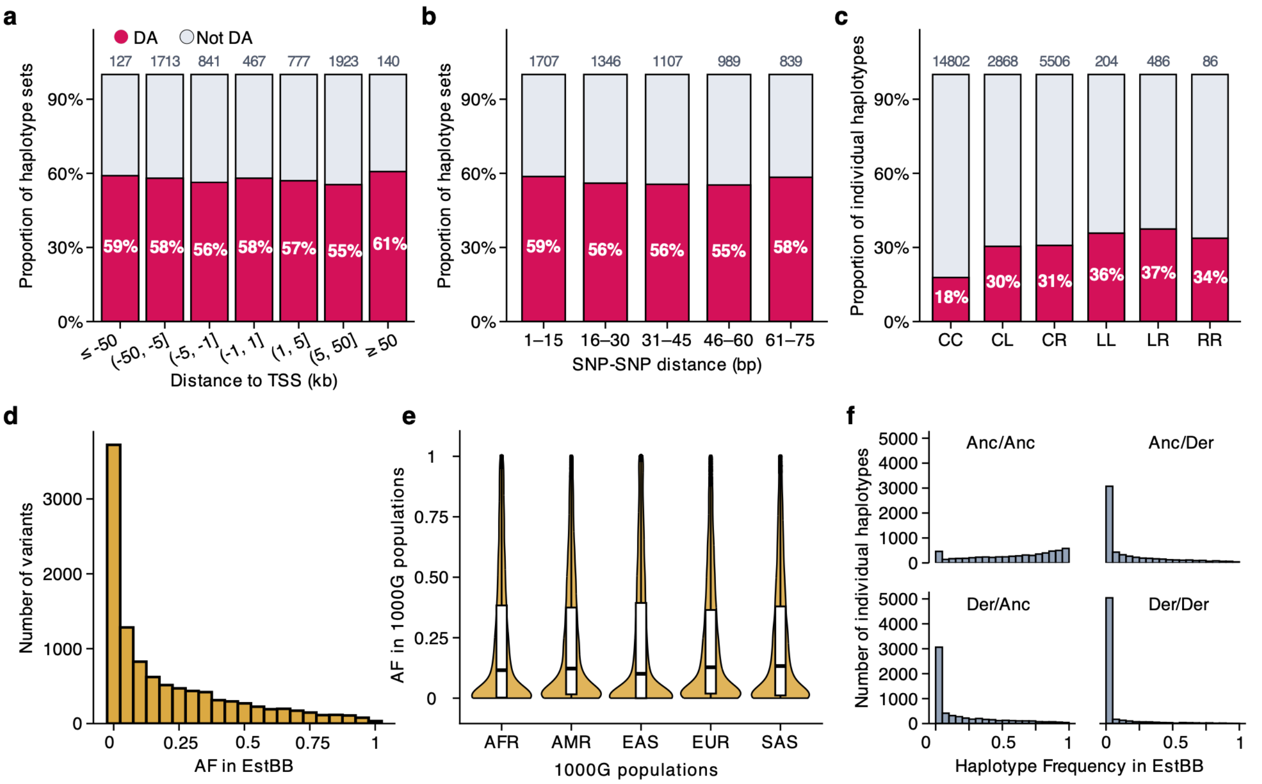


**Supplementary Fig. S7 Genomic and population characteristics of DA haplotype sets. a** Proportion of DA haplotype sets (red) across distance to TSS bins. Numbers on top of the bars show the total number of haplotype sets in the TSS bins. **b** Proportion of DA haplotype sets across SNV-SNV distance bins. **c** Proportion of individual DA haplotypes across AF combinations. CC is common/common; CL is common/low; CR is common/rare; LL is low/low; LR is low/rare; RR is rare/rare. **d** Allele frequency distribution of tested variants in EstBB data. **e** Allele frequency distribution of the tested variants in 1000G populations. **f** Haplotype frequencies in EstBB data stratified by ancestry combination.

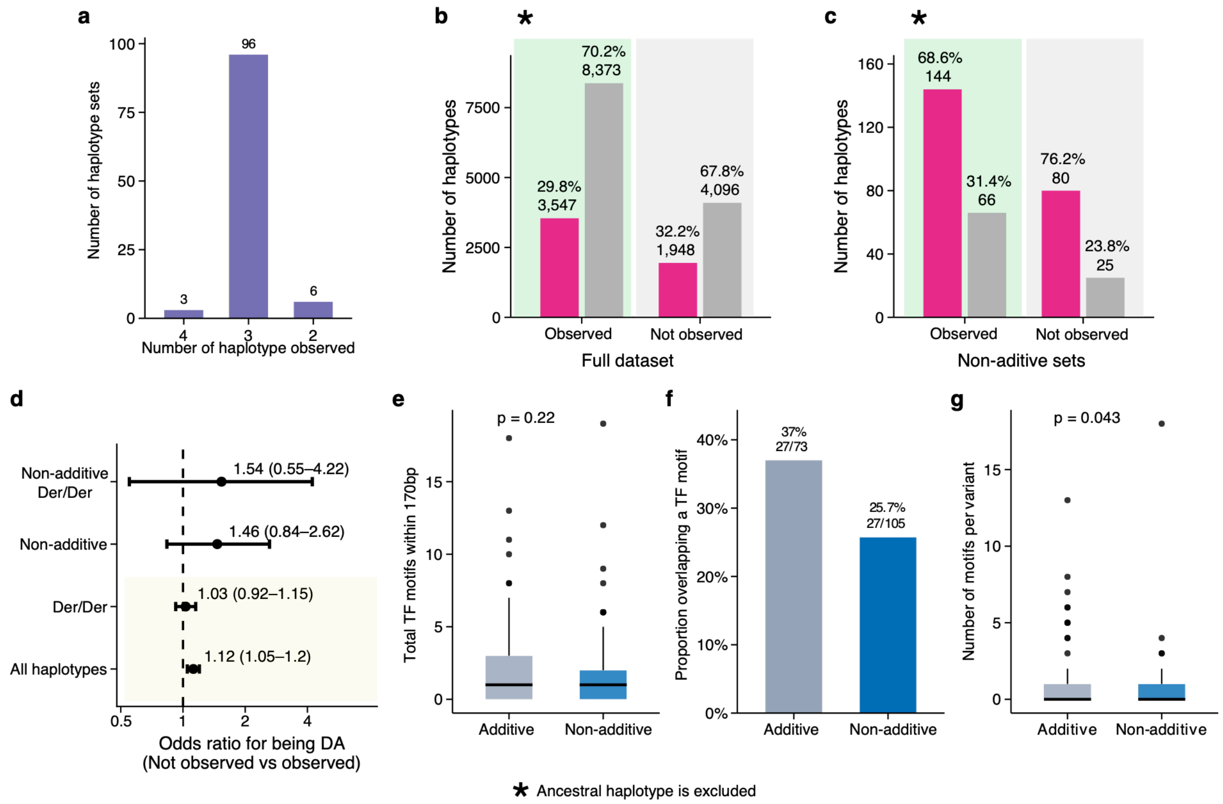


**Supplementary Fig. S8 Haplotype representation, DA, and TF motif characteristics**. **a** Number of haplotype sets with all 4 tested haplotypes, 3 and 2 haplotypes present in EstBB. **b** Number of observed vs not observed haplotypes in the full data, split by haplotypes that are DA (pink) and not DA (grey) in each group. **c** Number of observed vs not observed haplotypes in EstBB in the HC set, split by haplotypes that are DA and not DA. **d** Odds ratio of observed vs not observed haplotypes of being DA in whole data and in HC sets. Separated are Der/Der haplotypes for each category. **e** Distribution of known TF motifs found within 170 bp tested sequences in additive and non-additive sets. **f** Proportion of known TF motifs that directly overlapped at least 1 of the tested variants within additive and non-additive sets. **g** Distribution of unique TF motifs per variant within additive and non-additive sets.

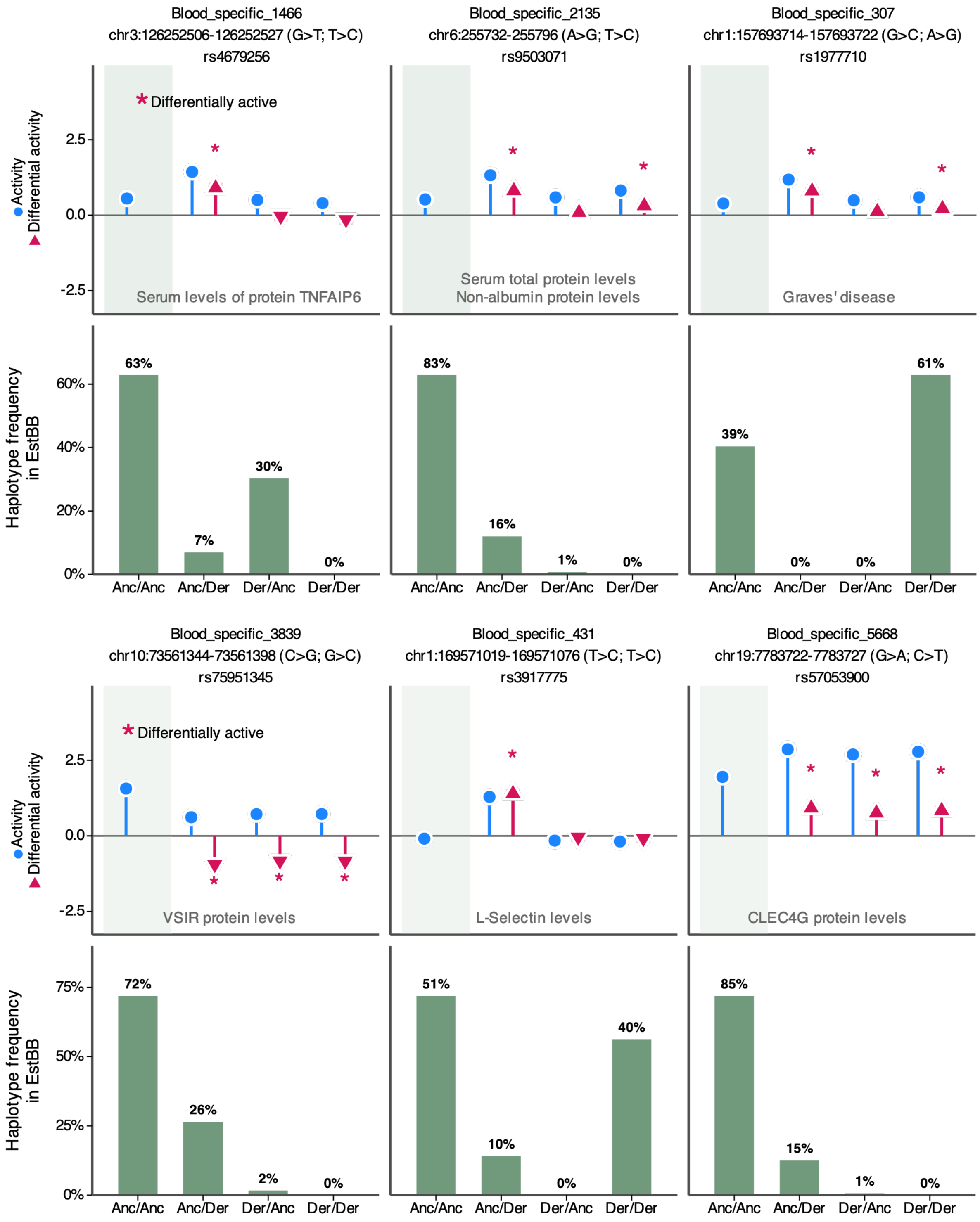

Blood_specific_1466
chr3:126252506-126252527 (G>T; T>C)
rs4679256
Blood_specific_2135
chr6:255732-255796 (A>G; T>C)
rs9503071
Blood_specific_307
chr1:157693714-157693722 (G>C; A>G)
rs1977710
* Differentially active
Activity
Differential activity
2.5
0.0
-2.5
Serum levels of protein TNFAIP6
Serum total protein levels
Non-albumin protein levels
Graves' disease
Haplotype frequency in EstBB
63%
7%
30%
0%
83%
16%
1%
0%
39%
0%
0%
61%
60%
40%
20%
0%
Anc/Anc
Anc/Der
Der/Anc
Der/Der
Blood_specific_3839
chr10:73561344-73561398 (C>G; G>C)
rs75951345
Blood_specific_431
chr1:169571019-169571076 (T>C; T>C)
rs3917775
Blood_specific_5668
chr19:7783722-7783727 (G>A; C>T)
rs57053900
VSIR protein levels
L-Selectin levels
CLEC4G protein levels
72%
26%
2%
0%
51%
10%
0%
40%
85%
15%
1%
0%
75%
50%
25%

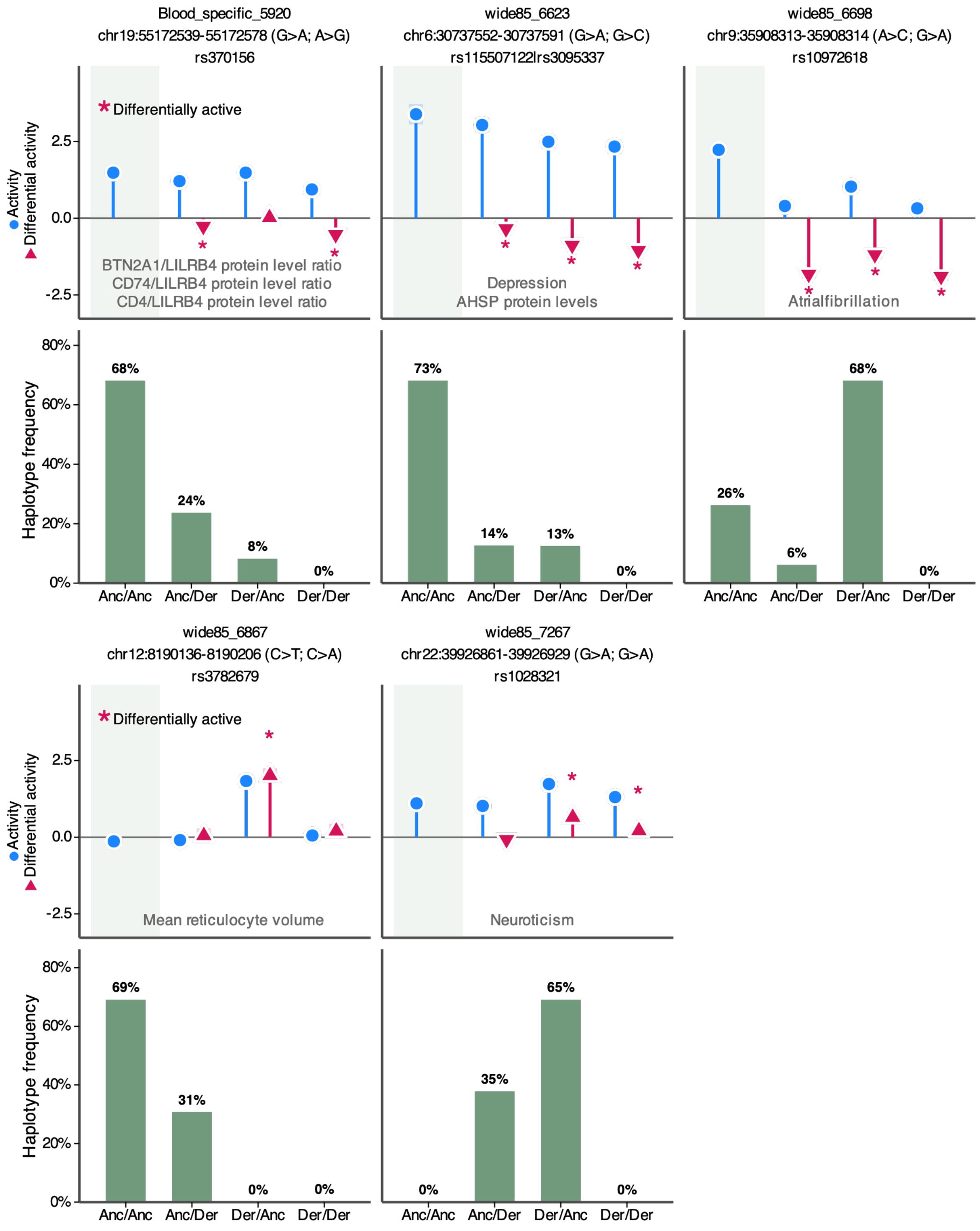


**Supplementary Fig. S9. Examples of non-additive haplotype sets with a known GWAS association.** Upper panels show haplotype activity and differential activity, with a GWAS disease association. Lower panels with green bars show haplotype frequencies in EstBB dataset.
